# Testing the two-step model of plant root microbiome acquisition under multiple plant species and soil sources

**DOI:** 10.1101/2020.03.11.988014

**Authors:** Hugo R. Barajas, Shamayim Martínez-Sánchez, Miguel F. Romero, Cristóbal Hernández-Álvarez, Luis Servín-González, Mariana Peimbert, Rocío Cruz-Ortega, Felipe García-Oliva, Luis D. Alcaraz

## Abstract

The two-step model for plant root microbiomes considers soil as the primary microbial source. Active selection of the plant’s bacterial inhabitants results in a biodiversity decrease towards roots. We collected *in situ* ruderal plant roots and their soils and used these soils as the main microbial input for single genotype tomatoes grown in a greenhouse. We massively sequenced the 16S rRNA and shotgun metagenomes of the soils, *in situ* plants, and tomato roots. Tomato roots did follow the two-step model, while ruderal plants did not. Ruderal plants and their soils are closer than tomato and its soil, based on protein comparisons. We calculated a metagenomic tomato root core of 51 bacterial genera and 2,762 proteins, which could be the basis for microbiome-oriented plant breeding programs. The tomato and ruderal metagenomic differences are probably due to plant domestication trade-offs, impacting plant-microbe interactions.

## Introduction

Soil and plant root-associated bacteria are relevant for plant health, which has already been noticed in the beginning of the 20^th^ century (Hiltner, 1904). It has been hypothesized that the microbiome could be related with crop quality (Hartmann *et al*., 2008). Soil is the most diverse microbial ecosystem, with up to 10^11^ bacterial cells per gram (Roesch *et al*., 2007). Soil properties such as pH, nutrient content, or moisture, and plant species can drive the soil microbiome composition (Fierer & Jackson, 2006; Lauber *et al*., 2009; Schlaeppi *et al*., 2014). Plants and soil interact at the rhizosphere, defined as the millimetric soil layer attached to plant roots. Plants play an active role in selecting their microbial inhabitants through root exudates, accounting from 5 to 20% of the photosynthetically fixed carbon and used by the microbes (Marschner *et al*., 2004). Plant-microbe interactions mainly occur in the rhizosphere (Berendsen *et al*., 2012). Some other known factors affecting the root microbial community structure are plant developmental stage (Inceoğlu *et al*., 2011), pathogen presence (Tian *et al*., 2015), and soil characteristics (Lundberg *et al*., 2012; Edwards *et al*., 2014). Plant-microbiome interaction has documented effects on plant growth and health; for example, the root microbiome composition has been associated with biomass increase in *Arabidopsis thaliana* (Swenson *et al*., 2000) and can also affect flowering time (Panke-Buisse *et al*., 2015).

A study of the *A. thaliana* microbiome using hundreds of plants and two soil sources concluded that root bacterial communities were strongly influenced by soil type (Lundberg *et al.,* 2012). Microbial diversity was reduced in the rhizosphere compared to the surrounding soil, suggesting that plants filter and recruit a microbiome subset; these observations have led to the two-step model of microbiome selection (Bulgarelli *et al.,* 2012; Bulgarelli *et al.,* 2013). This model considers soil abiotic properties in the soil microbiome (first step), and specific plant-derived rhizodeposits contribute to selecting differential microbes in the rhizosphere and the endosphere (second step) (Bulgarelli *et al.,* 2013). In the two-step model, α-diversity decreased in the following order: soils > rhizosphere > endosphere (Bulgarelli *et al*., 2013). However, a global-scale meta-analysis has reported that root microbiomes of multiple plant species (domesticated and wild) have a more substantial diversity than soils (Thompson *et al*., 2017).

This work explores the bacterial diversity by 16S rRNA gene massive amplicon sequencing and whole shotgun metagenomes to predict the protein diversity of 16 geochemically distinct Mexican soils, collected over a large geographical scale (Fig. 1A; Table 1). The collected soils were chosen based on country-wide edaphological charts (INEGI, 2014). We explored the role of soil in microbiome structuring of *in situ* ruderal plants, growing above the collected soils with multiple species and at several plant developmental stages. The collected soils were used as the substrate in a greenhouse experiment for growing tomatoes (*Solanum lycopersicum*), eliminating plant genotype variability as well as developmental, climatic, and watering variables. Finally, testing diverse soil groups allowed us to explore the tomato core root microbiome, which follows the two-step model for root microbiome selection. The ruderal plants do not follow the two-step model and have a larger diversity than their source soils.

**Figure 1.**
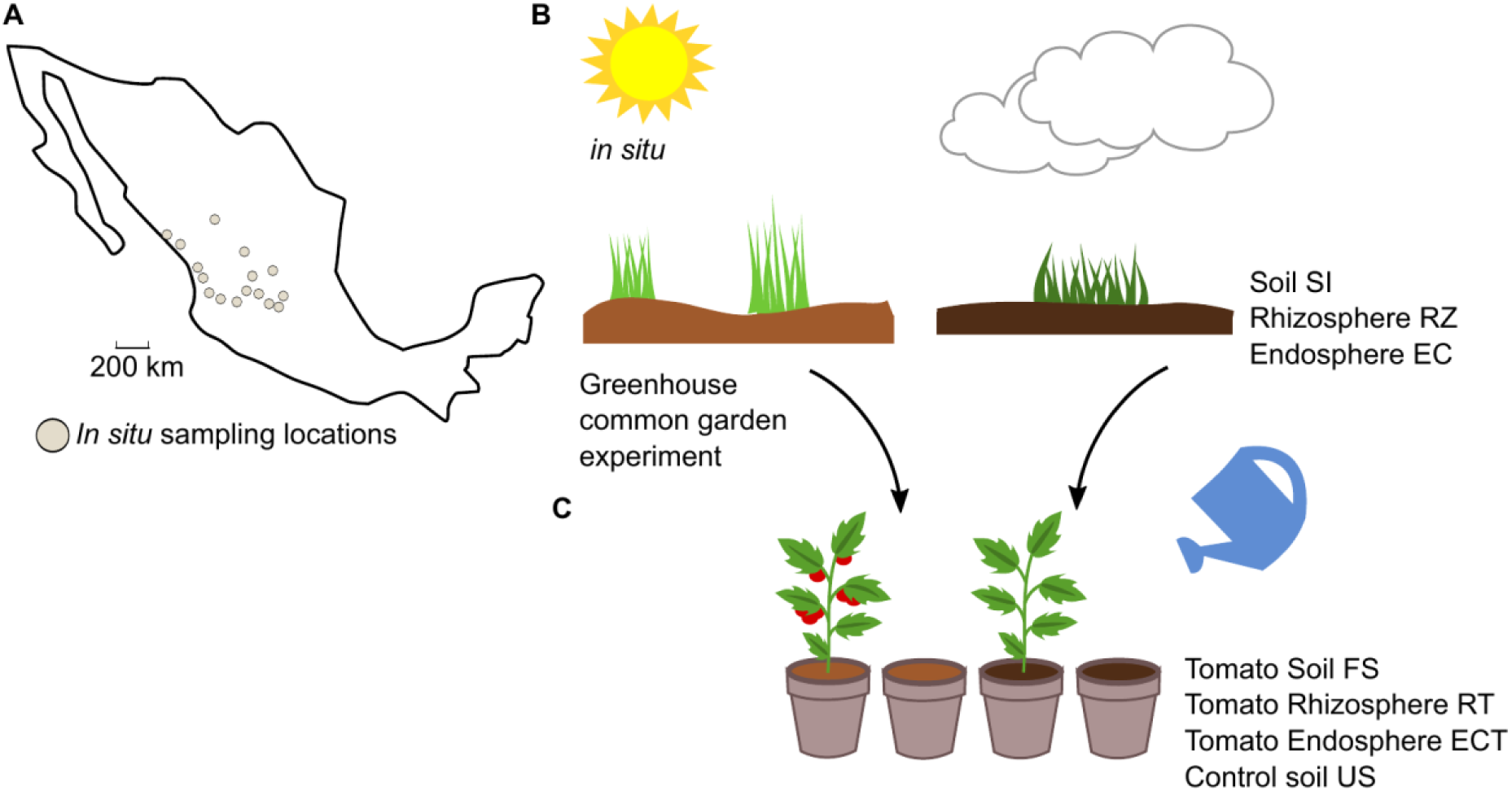
Experimental overview of the work. **A)** *In situ* sampling locations; sampling points were selected according to edaphological charts. **B)** The *in situ* plants were dependent on the weather and local environmental conditions, and we collected soil samples (SI) and roots of the dominant plant species in each locality. We extracted the rhizosphere (RZ) and endosphere (EC) metagenomic DNA. **C)** A common garden experiment was conducted in a greenhouse; the soil (SI) was used as a microbial inoculum to reduce environmental variability. Plant diversity was eliminated using tomato, with constant watering, and finally, we collected roots (RT), endosphere (ECT), final soil (FS), and control unplanted soil (US).

**Table 1.** Soil edaphological classification, abiotic properties, and tomato biomass production. Soils were classified according to the United Nations’ FAO edaphological charts. The main attributes of the soils used in this work are sampling location, altitude (m.a.s.l.), Lang’s Aridity Index (AI), and contents of carbon, nitrogen, and phosphorus. Initial refers to source soil (SI) and final to common garden experiment output soil (FS) measurements.

| Site | Soil group | pH<br>(SI) | pH<br>(SF) | Total<br>N<br>(mg/g)<br>(SI) | Total<br>N<br>(mg/g)<br>(SF) | Total<br>C<br>(mg/g)<br>(SI) | Total<br>C<br>(mg/g)<br>(SF) | Total<br>P<br>(mg/g)<br>(SI) | Total<br>P<br>(mg/g)<br>(SF) | Tomato<br>biomass<br>(SF)<br>(g) | AI | Altitude | Latitude | Longitude | Collection<br>date |
| --- | --- | --- | --- | --- | --- | --- | --- | --- | --- | --- | --- | --- | --- | --- | --- |
| JAL4 | Phaozem | 7.2 | 6.0 | 0.623 | 11.721 | 16.503 | 31.003 | 0.009 | 0.16 | 0.221 | 48.8 | 1490 | 20.849 | -103.855 | 11/8/2014 |
| NAY2 | Cambisol | 5.7 | 6.1 | 6.83 | 16.667 | 19.347 | 22.086 | 0.453 | 0.284 | 0.083 | 56.5 | 38 | 21.836 | -105.083 | 11/8/2014 |
| JAL3 | Luvisol | 6.5 | 6.8 | 3.99 | 17.799 | 32.863 | 25.683 | 0.475 | 0.263 | 0.090 | 46.1 | 1862 | 20.821 | -102.798 | 11/6/2014 |
| NAY3 | Cambisol | 6.9 | 6.9 | 2.027 | 11.584 | 9.924 | 13.569 | 0.157 | 0.197 | 0.204 | 47.1 | 4 | 22.142 | -105.261 | 11/8/2014 |
| SLP1 | Kastanozem | 6.9 | 7.8 | 9.702 | 5.898 | 79.348 | 146.37 | 0.447 | 0.287 | 0.767 | 22.1 | 2130 | 21.988 | -101.277 | 11/10/2014 |
| ZAC1 | Cambisol | 5.0 | 7.2 | 6.433 | 12.961 | 4.921 | 5.117 | 0.324 | 0.347 | 0.017 | 25 | 2221 | 22.818 | -102.696 | 11/10/2014 |
| SIN1 | Phaozem | 7.3 | 7.4 | 7.523 | 2.095 | 16.862 | 24.452 | 0.377 | 0.42 | 0.223 | 29.8 | 23 | 22.988 | -105.880 | 11/8/2014 |
| JAL5 | Lithosol | 7.4 | 7.9 | 2.158 | 24.011 | 5.067 | 5.427 | 0.235 | 0.192 | 0.199 | 44 | 1328 | 21.179 | -104.540 | 11/8/2014 |
| JAL1 | Planosol | 7.5 | 7.6 | 1.892 | 1.339 | 9.482 | 14.425 | 0.118 | 0.164 | 0.313 | 32.9 | 1810 | 21.162 | -101.853 | 11/6/2014 |
| AGS1 | Planosol | 8.0 | 8.3 | 5.671 | 12.46 | 19.616 | 22.01 | 0.268 | 0.28 | 0.269 | 29.2 | 2059 | 21.828 | -102.121 | 11/10/2014 |
| GTO1 | Vertisol | 8.3 | 8.6 | 12.298 | 13.387 | 22.733 | 18.546 | 0.378 | 0.323 | 0.259 | 32.9 | 1708 | 20.580 | -100.947 | 11/11/2014 |
| GTO3 | Vertisol | 8.4 | 8.4 | 2.624 | 6.008 | 10.559 | 10.378 | 0.271 | 0.323 | 0.180 | 29.2 | 2012 | 20.897 | -100.674 | 11/11/2014 |
| JAL2 | Phaozem | 8.6 | 9.0 | 2.075 | 8.646 | 14.929 | 13.285 | 0.01 | 0.336 | 0.219 | 38.2 | 1791 | 21.253 | -102.298 | 11/6/2014 |
| GTO2 | Vertisol | 8.6 | 8.7 | 7.089 | 13.948 | 18.038 | 17.722 | 0.4 | 0.417 | 0.156 | 33.8 | 1710 | 20.721 | -101.329 | 11/11/2014 |
| DGO1 | Planosol | 8.9 | 8.9 | 5.018 | 10.838 | 23.36 | 23.773 | 0.251 | 0.248 | 0.156 | 33.8 | 1906 | 24.006 | -104.385 | 9/11/2014 |
| SIN2 | Regosol | 9.1 | 8.8 | 18.578 | 4.91 | 49.131 | 50.98 | 1.103 | 0.157 | 0.314 | 29 | 10 | 23.293 | -106.479 | 9/11/2014 |

## Results

### Soil geochemical description

Total nutrient concentration (C, N, P), pH, and Lang’s aridity index were calculated and considered as soil abiotic properties (Table 1). We observed an increase in N concentrations in 12/16 samples, while total C increased in 11/16 samples and P decreased in 7/16 samples (Table 1). Tomatoes planted in SLP1 and SIN2 exhibited a reduction in their total N concentrations; in SLP1, this is explained as plant biomass generation, and in SIN2, a coastal dune N was probably drained through watering (Table 1, Fig. S2). Only two samples changed their pH profiles (Table 1). Ordination analysis showed clustering apart of source soils (SI) from final greenhouse soils (FS) and evidenced the modifications derived from the common garden experiment (Fig. S2A; Table 1).

### Microbiome diversity in the source soils, ruderal plants, and tomatoes

A total of 106 amplicon libraries (16S rRNA gene V3-V4) were sequenced (Fig. 1, Supplementary Table S1). After quality control and assembly, 5,211,969 sequences were recovered. Subsequently 2,570,541 operational taxonomic units (OTUs; 97% identity) were clustered. After discarding singleton, mitochondrial, chloroplast, chimeras, and non-matching sequences, a total of 271,940 OTUs were the base for further analysis. The average *in situ* source soil (SI) OTU number (SI = 2,143) was lower than that in ruderal plants rhizospheres (RZ) and endosphere (EC) (RZ = 18,158; EC = 19,885 OTUs). Common garden soil (FS) and controls (US) had similar OTU averages (FS = 3,084; US = 2,882). Tomato rhizosphere (RT) and endosphere (ECT) samples had a higher OTU average than the SI, but a smaller average compared to FS (RT = 2,474 OTUs, EC = 2,088) (Supplementary Table S2).

We found 586 shared bacterial genera between soils and roots (rhizosphere and endosphere) of tomatoes and ruderal plants (Fig. 2A). The source soils had 8 unique genera and shared most (98.78%) of their microbes with tomatoes or ruderal plants. The largest amount of unique genera (46.21%) was found for the ruderal plants, sharing the most bacterial genera with the tomato and the soils (53.78%). The tomato root microbiomes had 14 unique bacterial genera (1.9%), 4 were only shared with soils (0.53%), while most genera were shared with soils and ruderal plants (97.53%). Another comparison showed that the tomato core microbiome had 51 bacterial genera, while ruderal plants and core soils had 87 and 16 bacterial genera, respectively. Cores were defined as detected genera in all of the sample types compared (RT, soils, and RZ). Complete information on unique and shared OTUs is available (Supplementary Table S3).

**Figure 2.**
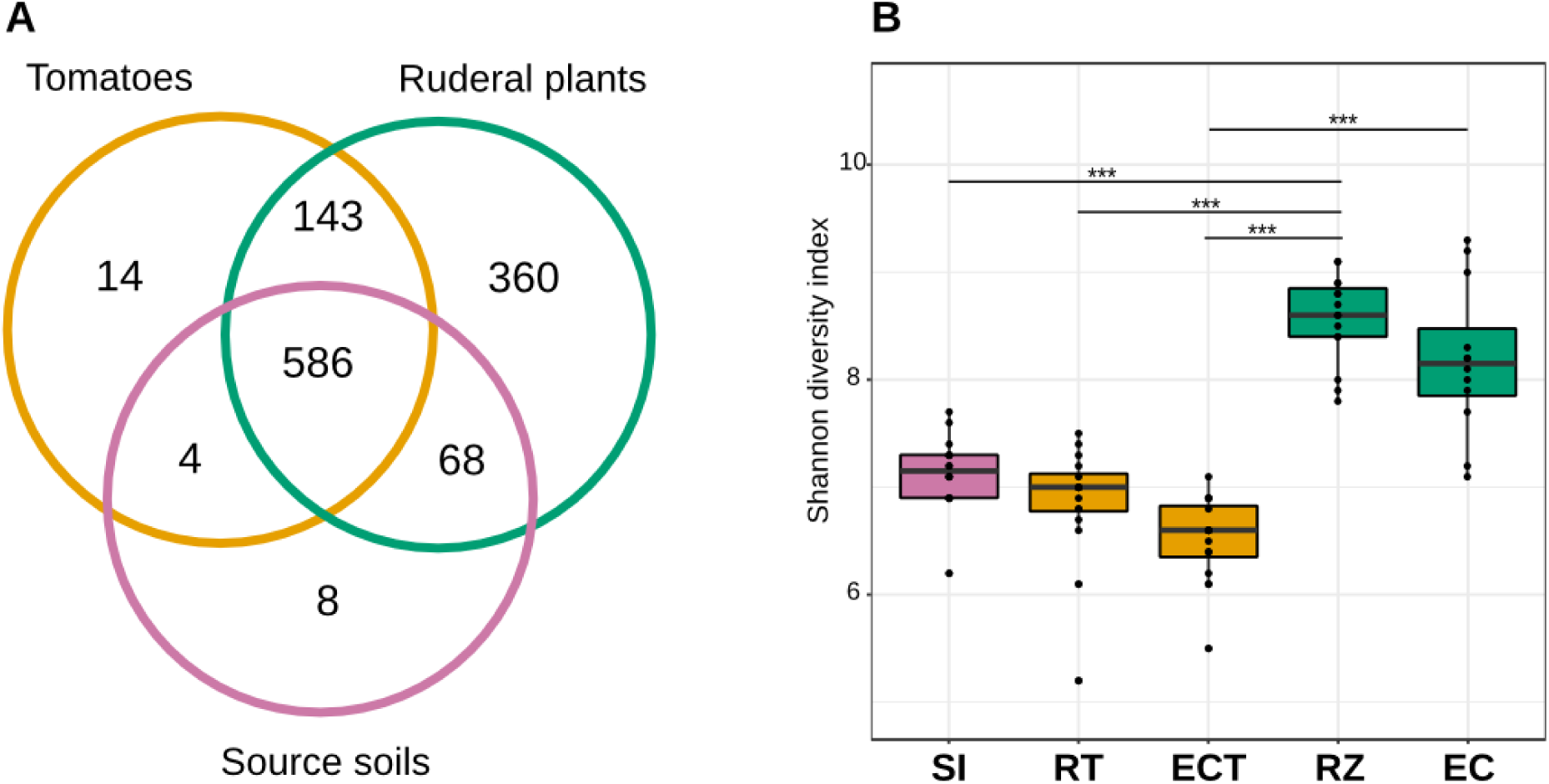
Alpha diversity and richness of soil, rhizosphere, and endosphere of tomato and ruderal plants. **A)** Venn diagram showing the number of shared bacterial genera in roots (endosphere + rhizosphere) and soils. **B)** Boxplots showing the OTU Shannon diversity index (H’) of source soils (SI), tomato rhizosphere (RT), tomato endosphere (ECT), *in situ* plants rhizosphere (RZ), and *in situ* ruderal plant endosphere (EC).

We analyzed the α-diversity of soils, rhizospheres, and endosphere microbiomes through the Shannon diversity index (H’). The H’ index values show that soils were more diverse (H’ = 6.1 to 7.6) than the tomato rhizosphere (H’ = 5.2 to 7.4) or the tomato endosphere (H’ = 5.5 to 7.1), thus fitting the two-step model for microbiome selection. However, when comparing the soil to the ruderal plant root microbiomes, higher H’ values were observed in the rhizosphere (H’ = 7.4 to 9.1) and even in their endosphere microbiomes (H’ = 7.0 to 9.2) compared to their soils (H’ = 6.1 to 7.6), not adjusting to the two-step model (Fig. 2B, S3, Supplementary Table S2).

We performed β-diversity analysis based on the weighted UniFrac community distance matrix to dissect the role of soil in the establishment and structure of rhizo- and endosphere microbiomes in both ruderal and *S. lycopersicum* plants (Fig. 3). The weighted UniFrac dendrogram grouped the samples into three major clusters: Cluster (I) contains only tomato-associated microbiomes, cluster (II) includes soil and ruderal plant microbiomes, and a mixed cluster (III) includes soil, tomato, and ruderal plant microbiomes (Fig. 3). The clustering of the three groups is supported by ANOSIM (R = 0.7257; *p* < 0.001; 999 permutations).

**Figure 3.**
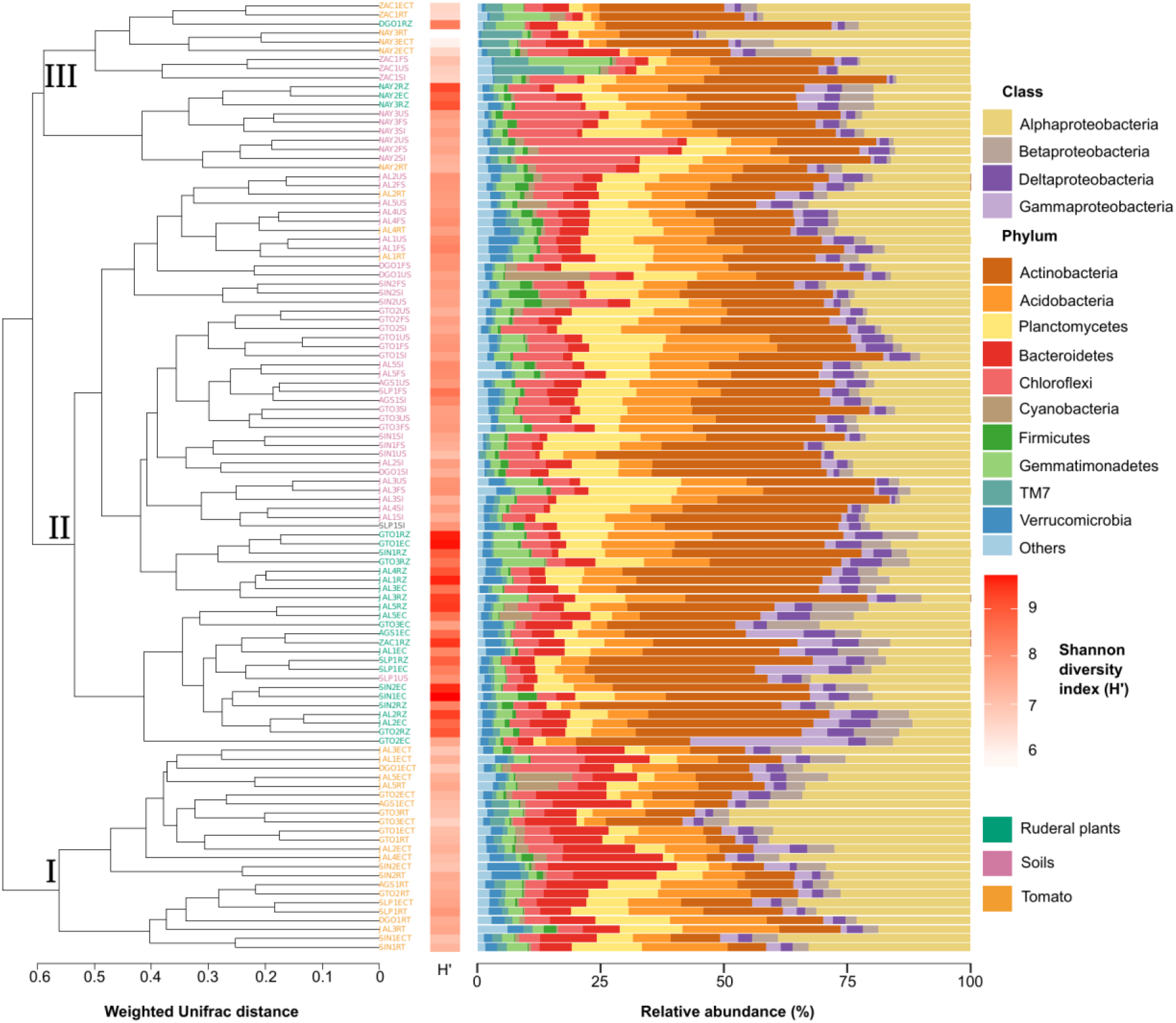
Host genotype and soil influence on microbial community structure (16S rRNA gene). On the left, a weighted Unifrac dendrogram shows β-diversity and phylogenetic similarity between soil, tomato, and ruderal plants. Each location is indicated at the dendrogram terminal nodes with a three-letter key for sampling location and suffix indicating type: initial source soil (SI), final soil (FS), unplanted soil (US), tomato rhizosphere (RT), tomato endosphere (ECT), ruderal plant rhizosphere (RZ), and ruderal plant endosphere (EC). Phyla diversity (H’) in each sampled microbiome is shown as a horizontal heatmap. Bar plots show bacterial phyla relative abundance in each sample.

Pairwise distance values were calculated between every sample in the weighted UniFrac dendrogram to evaluate the distance patterns and cohesion found inside and between each of the described clusters. The average internal distances were 0.5041 for cluster (1), 0.5058 for cluster (2), and 0.4787 for cluster (III). The measured distance between any terminal node of cluster I against any tip in either cluster (II) or (III) was 0.6608. The tomato-associated cluster (Fig. 3. cluster I) grouped 10/16 tomato rhizospheres, along with 13/16 of the tomato endospheres, suggesting a plant genotype-dependent role in root microbiome establishment.

The tomato and ruderal samples that were not clustered with their own were clustered closer to their source soils, remarkably acid soils, indicating pH properties as microbiome structuring factor (Table 1).

*In situ* samples of soils and ruderal plants were dominated by Actinobacteria, with a significantly (ANOVA *p* < 2e-16) lower abundance in tomato roots (Fig. 3; Fig. S4; Supplementary Table S5). Proteobacteria were significantly enriched in tomatoes (ANOVA *p =* 1.82e-15) when compared to soils and ruderal plants. The class α-Proteobacteria was the most abundant one in tomatoes, with significant enrichment (ANOVA *p* < 2e-16) when compared to ruderal plants and soils. The β, γ, and δ-Proteobacteria were more abundant in ruderal plants (*p* < 0.05) than in tomatoes and soils. Bacteroidetes were enriched in tomato roots (ANOVA *p <* 1.34e-15) when compared to soils and ruderal plants (Fig. 3; Fig. S4; Supplementary Table S5).

### Proteobacteria are shown at the class level in the bar plots

We used DESeq2 to compare and identify significantly (*p* < 0.01) enriched OTUs (Supplementary Table S6). We found five differential OTUs assigned as *Sphingobium, Caulobacter, Asticcacaulis, Arthrospira,* and *Kaistobacter* in the tomato rhizospheres compared to their source soils (Fig. S5A). In contrast, in the endospheres, we found 12 enriched OTUs belonging to the same genera present in the rhizosphere, along with *Agrobacterium* and *Lacibacter* (Fig. S5B). The comparison between ruderal plant roots and soils showed 21 differentially abundant OTUs in the rhizospheres (Fig. S5C) and 29 enriched OTUs in the endosphere (Fig. S5D), most belonging to Actinobacteria (Supplementary Table S6). Additionally, we compared the sets of soils and their controls (Fig. 1) and did not find any shared OTUs whose abundance differed significantly.

### Shotgun metagenomic diversity in source soils, ruderal plant rhizospheres, and tomato rhizospheres

We sequenced 50.1 Gb in a total of 15 SI, RT, and RZ metagenomes. After quality control, we obtained 464,372,598 high-quality paired-end reads (μ = 27,316,035 ± 2,943,233 per sample), which were used as input to an assembly that yielded 12,677,118 contigs (μ = 745,713 ± 366,001 per sample), with an average N50 of 176 ± 51 bp and the longest contig average length of 45,645 bp. Subsequently, we were able to compute a total of 12,272,971 predicted peptides (μ = 708,835 ± 332,770 per sample) (Supplementary Table S7). Protein redundancy was reduced using proxy-genes of matches to known proteins and protein-clustering alignments (70% identity). After clustering and matching, protein annotation was performed using the M5NR database (see Methods), resulting in 3,147,929 proteins; only 411,432 (13.07%) were annotated based on homology against the M5NR database.

We compared the shared set of proteins between soils, ruderals, and tomatoes, resulting in a set of 43,305 proteins detected at least once for every sample type (Fig. 4A). Most of the union set proteins (93%) were annotated. Tomatoes shared with the soils 8.49% of their predicted proteins, while ruderal plants shared 8.72% of the identified peptides with the soil. Tomatoes shared more coding genes with ruderal plants (8.85%) than with soil (8.49%) (Fig. 4A). Different sets of proteins for each sample showed the largest novelty in soil (88.83%), followed by ruderal plants (87.46%) and tomatoes (86.36%) (Fig. 4A). Although the largest number of unique proteins could be the result of an enthusiastic computer prediction, it was interesting that the tomato had the largest amount of annotated proteins (12.10%) compared to ruderals (9.97%) and soils (6.75%), maybe reflecting the larger previous genomic information in agricultural microbes, being scarcer in wild plants, and the soil microbes (Fig. 4A). Complete lists of identified proteins are available as supplementary material (Supplementary Table S8).

**Figure 4.**
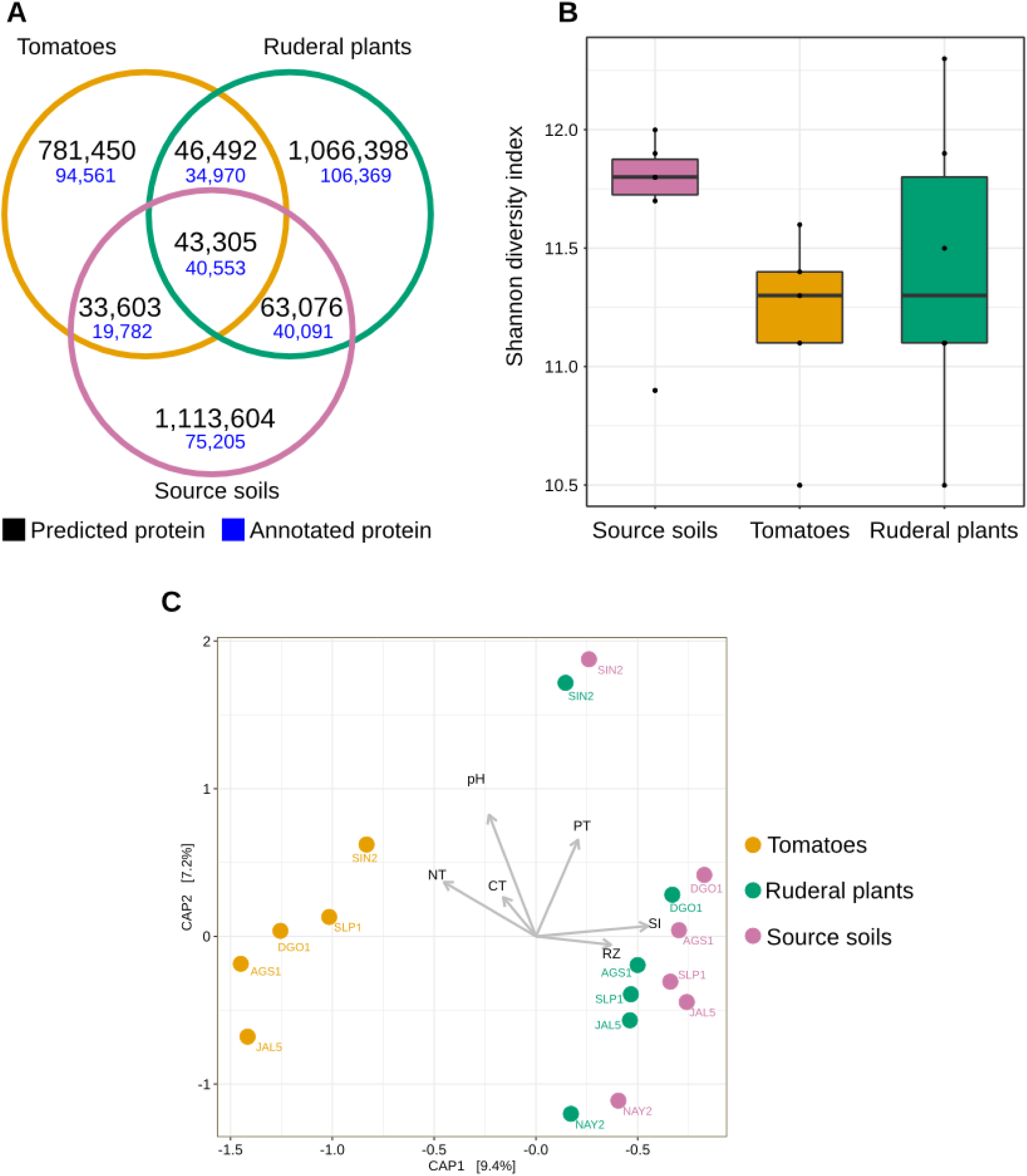
Shotgun metagenomics diversity of soil and rhizosphere microbial communities. **A)** Venn diagram showing the number of shared and unique annotated protein families (70% identity) in soil and rhizosphere. **B)** Boxplots showing the Shannon diversity index based on the total number of predicted proteins in soil and rhizosphere. **C)** Constrained analysis of principal coordinates (CAP), calculated from the total number of predicted proteins for all sequenced soil and rhizosphere metagenomes using Bray-Curtis dissimilarity. Vectors display the environmental factors: CT = Total Carbon concentration, NT = Total Nitrogen concentration, PT = Total Phosphorus concentration, RZ = ruderal plant rhizosphere, SI = Initial soil.

We compared the protein α-diversity using the Shannon diversity index (H’) based on the total number of predicted proteins (Supplementary Table S9). Soil diversity had a higher median (H’ = 11.8) than tomato diversity (H’ = 11.3) and ruderal plant diversity (H’ = 11.3), without significant differences (Fig. 4B, S6). To test the hypothesis that the tomato predicted metaproteome is divergent from those of the soil and ruderal plants, as suggested by the 16S β-diversity dendrogram (Fig. 3), we performed a constrained analysis of principal coordinates (CAP) ordination (Fig. 4C). We used the protein abundance as CAP input and constrained the analysis by sample type, pH, total N, C, and P. The CAP explained 16.6% of the total observed variance, with CAP1 (9.4%) splitting the tomatoes from ruderals and source soils. Ruderal plants were closer to the soils, but were not mixed with them (CAP, Bray-Curtis distance, PERMANOVA 9,999 permutations, *p* < 1e-4). The split tomato and ruderal-soil groups are also supported by ANOSIM (R = 0.4568; *p* < 0.001; 999 permutations) (Fig. 4C). Correlations with the measured geochemical variables with the two CAP axes showed positive correlations of P, source soils (SI), and ruderal plants (RZ), while negative correlations were observed for pH, N, and C (Fig. 4C).

We were able to bin and classify 38% ± 1.66 of the sequenced metagenomic reads to multiple taxa. Bacteria accounted for 88.14% of the identified matches, followed by 7.09% of Eukaryota, with almost half of the hits corresponding to Fungi (3.83%) (Supplementary Table S10). Further work will explore eukaryotic, archaeal, and viral diversity of the sequenced metagenomes. In soils, the most abundant bacterial species binned was *Solirubrobacter soli* (Actinobacteria), which was also highly abundant in ruderals and tomatoes. *Sphingomonas* sp. URHD0057 (α-Proteobacteria) were most enriched in ruderals, along with *S. soli* and the Rhizobiales *Rhodoplanes* sp. Z2-YC6860. The 16S data showed that Bacteroidetes were significantly enriched in tomato roots compared to soil and ruderals; metagenomic bins confirm the 16S rRNA gene trends. Within the principal metagenomic bins, we found *Ohtaekwangia koreensis*, *Flavobacterium terrae, Niastella vici, Chryseolinea serpens,* and the metagenome-assembled genome of a *Chitinophagaceae* bacterium IBVUCB2 as Bacteroidetes species.

We analyzed a functional summary of the sequenced metagenomes using SEED subsystem gene ontology (Fig. 5). The largest category was described as clustering-based subsystems, which include protein families quite diverse from the CRISPR, sugar metabolism, other known categories, as well as hypothetical proteins. We only found small differences (Tukeýs HSD) in iron acquisition metabolism (*p* = 0.07), cell wall and capsule genes (*p* = 0.06) between soils and tomatoes. Significant differences (*p* = 0.017) were found between ruderals and tomatoes in sulfur metabolism genes (Fig. 5).

**Fig. 5.**
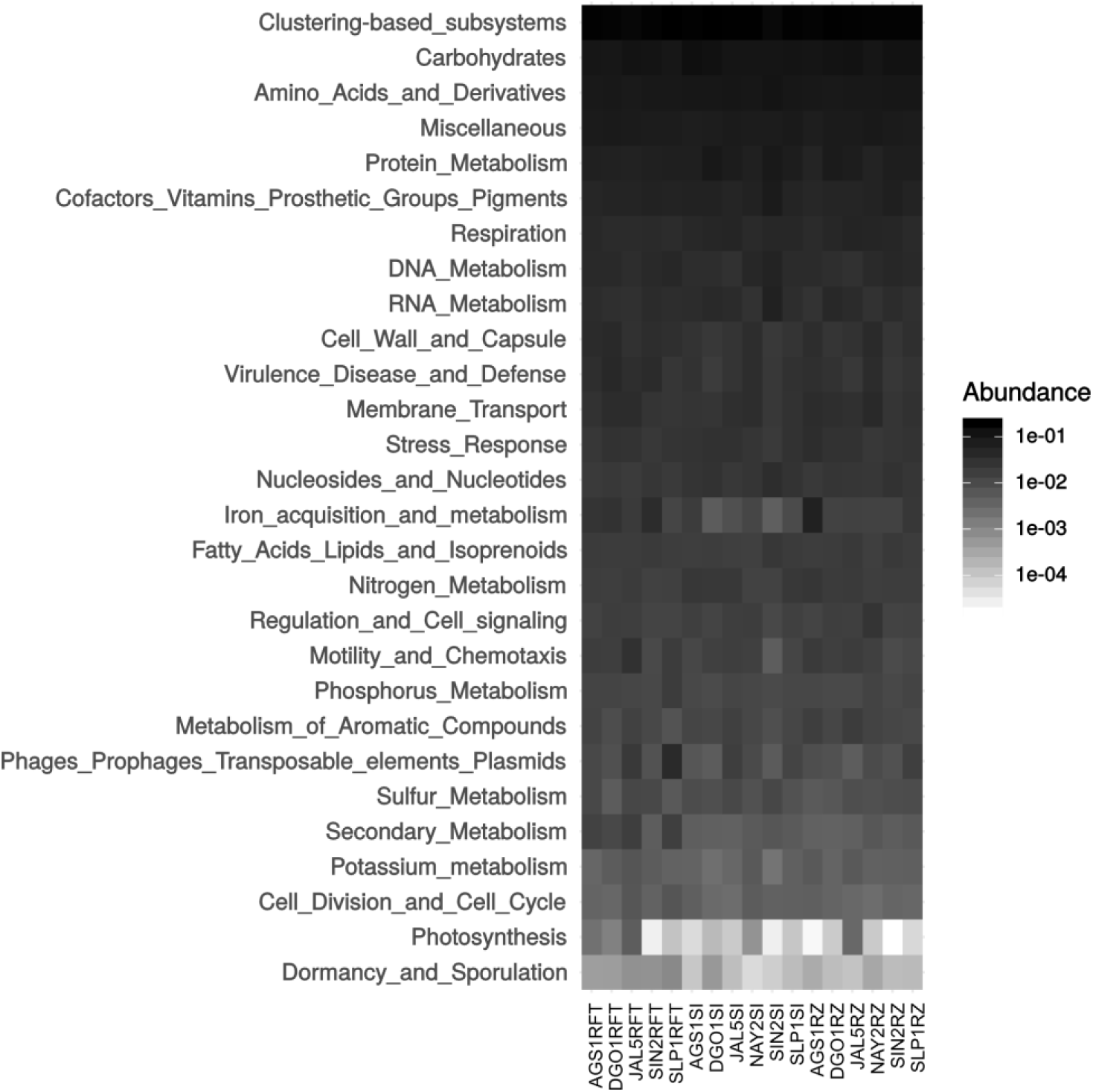
Summary of metagenomic functional profiles. Heatmap describing the level 1 SEED subsystems ontology annotations in each row. Although only using the 138,627 M5NR matches to the SEED, representing 4.4% of the total dataset, it was helpful to describe the main molecular functions. Columns represent each metagenome, and the first four alphanumeric codes are for location; suffixes indicate sample type: RFT are tomato roots, SI is source soil, and RZ represents ruderal plant metagenome.

### Enriched proteins in the rhizospheres-soil comparison

Pairwise comparisons were made using DESeq2 to find significant (*p* < 0.001) enrichments. Comparing tomatoes (RT) and soils (SI), we identified 67 enriched proteins in RT involved in motility, chemotaxis, and biofilm formation (*e.g.,* LuxR, CheY, diguanylate cyclase), complex carbohydrate degradation (*e.g.,* xyloglucanase, cellulase Cel5F), antibiotic resistance (*e.g.,* β-lactamase class C), iron metabolism (*e.g.,* TonB), and sporulation (*e.g.,* SpoIIIE), as well as secretion system-related proteins (*e.g.*, exo-sortase) (Fig. S7).

When comparing ruderals (RZ) and tomatoes (RT), we found 16 enriched proteins in RT and 11 in RZ (Fig. S8). The lowest number of enriched proteins between tomatoes and ruderals indicates that their shared set contains common features in plant-microbe interactions. The RZ-enriched proteins, compared to RT, included transporters and, interestingly, osmotic sensor components (*e.g.,* osmosensitive K channel histidine kinase). The RT-enriched proteins, compared to RZ, included several peptidases (*e.g.,* M17 leucyl aminopeptidase) and some horizontal gene transfer elements (*e.g.,* integrase-recombinase, ISRSO17 transposase, bacteriophage N4 adsorption protein B). Interestingly, multiple similar proteins enriched in the RT-SI comparison were also enriched in the RT-RZ (*e.g.,* β-lactamase class C, glycoside hydrolases), remarking the host genotype filtering of RT. Finally, comparing RZ-SI only showed two RZ-enriched proteins, indicating the similarities between soil and ruderals (Fig. S9). The full list of overrepresented proteins for each comparison is available (Supplementary Table S11).

### The tomato rhizosphere, soil, and ruderal plant core metagenomes

It seems that the tomato was highly selective about its microbial inhabitants; we found 2,762 protein families ubiquitous in all tomato roots tested (Fig. S10). We used the protein annotation to reduce the dataset to 1,777 core proteins and to only 1,353 exclusively in tomato (Supplementary Table S12). The core tomato metagenome was contrasting to the soil with only 162/639 and the ruderal metagenome with just 143/694 core-exclusive proteins.

Some essential proteins were expected to be part of the core metagenomes and worked as controls for our searches, such as ribosomal proteins, DNA and RNA polymerases, gyrases, chaperonin GroEL, and we found them all within the tomato core metagenome. This metagenome contained nitrogen regulation genes via denitrification (*nosZ*) and nitrate reductase genes (*nasA, nirB, nrfA*) to obtain ammonia. Glutamate, glutamine synthetases, and their transferases were also detected in the RT core metagenome and could be regulating both amino acid synthesis and ammonia. Additional nitrogen storage proteins were detected, such as cyanophycin synthetase (CphA) and cyanophycinase, within the RT metagenomic core; cyanophycin is a non-ribosomal peptide built by aspartic acid and arginine. Further, we found allantoinase and allantoate amidohydrolase genes, which are responsible for allantoin degradation to ammonia. Other tomato core proteins were patatin-like phospholipase proteins. While significantly enriched in ruderals, leucyl aminopeptidase was also ubiquitous in tomato metagenomes. The complete M5NR identifiers and core metagenomes are available (Supplementary Table S13).

## Discussion

With the common garden experiment, we increased biological activity, reflected in the soil N and C overall increases of the soils after the experiment, and the changes in bacterial diversity (Table 1). The high abundance of Actinobacteria in the source soils and the switch to a Proteobacteria dominance in the FS suggests processes such as biological nitrogen fixation and microbial biomass increments. Both Actinobacteria and Proteobacteria are capable of nitrogen fixation since their genomes contain nitrogenases (Boyd & Peters, 2013). However, it seems that plants such as tomatoes as other agricultural species favor Proteobacteria, while ruderals and soils depend upon Actinobacteria. Another possible explanation for the soil C increase is carbon deposition by plant root exudates (Canarini *et al*., 2019). We observed that the microbiome (16S rRNA gene) distribution was largely driven by the host interaction.

Most of the tomato samples were closer to each other than to their source soils (Fig. 3). The ruderal plants were mostly clustered together, with the highest observed diversity (H’; Fig. 3) and as a sister clade to their source soils. The explored abiotic differences showed that soil pH was a good predictor for microbiome distribution, mostly for acidic soils (Cluster III, Fig. 3, Table 1). The pH as a microbiome predictor has been reported before (Fierer & Jackson, 2006; Mannisto *et al*., 2007).

Different studies have confirmed the two-step model of root microbiomes (Lundberg *et al*., 2012; Bulgarelli *et al*., 2013, Edwards *et al*., 2015). Here, we found that taxa diversity of tomato roots is lower than the diversity found in the surrounding soil, thus consistent with the two-step model (Bulgarelli *et al*. 2013). The overall diversity decrease in the tomato roots is consistent with the enrichment of certain bacterial groups capable of close plant interactions through specific molecular mechanisms (*e.g.,* chemotaxis responsive, plant degradation enzymes) (Bais *et al*, 2006; Compant *et al*., 2010). The higher taxa diversity observed in ruderals is opposed to the two-step model. However, previous reports show a larger diversity in rhizospheres than in soils comparing different biomes (Thompson *et al*., 2017). Each plant can attract and select specific microorganisms depending on plant-genotype-dependent chemical formulation of rhizodeposits and cell wall features, resulting in specificity for microbiome selection (Fitzpatrick *et al*., 2018; Schlaeppi *et al*., 2014; Bulgarelli *et al*., 2013).

Some considerations must be made for ruderal plants since they are present in environments where they are not the only plant species but part of a plant community that could be broadening the rhizosphere effect. Different plant species or genotypes, as well as plant age, have been reported to attract specific bacterial communities (Marschner *et al*., 2004; Micallef *et al*., 2009; Baudoin *et al*., 2002). Additionally, natural variation in the climatic conditions was site-specific, while in the common garden experiment, tomato plants were watered regularly and climatic variation was minimized. The increased abundance of Actinobacteria in both soils and ruderal plants can also be a product of environmental water limitations, which directly affects the proportions of these phyla in arid soils, while humid sites usually have larger Proteobacteria abundances (Neilson *et al*., 2017). Proteobacteria have faster duplication times than Actinobacteria (Ramin & Allison, 2019). Most of the ruderals in our study were grasses (Supplementary Fig. S1), and recently it was reported that grasses rhizospheres (Poaceae family) were enriched in Actinobacteria under drought conditions (Naylor *et al*., 2017). Ruderal plants might have a larger α-diversity because of the rhizosphere micro-environmental conditions, analogous to an oasis in the dry soil.

Additionally, the number of plant species growing in the soil affects belowground microbial community diversity, biomass, and respiration rates, thereby impacting plant diversity (Wu *et al*., 2019). The large abundance of Actinobacteria has practical explanations in plant interactions; they have been used as biocontrol agents isolated from soil and rhizospheres, and they are secondary metabolite producers such as antibiotics or plant growth-promoting molecules such as indole acetic acid (El-Tarabily *et al*., 2010; Brader *et al*., 2014, Sreevidya *et al*., 2016).

By using the tomato (fixed plant genotype), we imposed a selective factor, since the plant-derived chemotactic signals and photosynthates should be similar, independent of the soil.

There are reports about enrichments of specific bacterial groups, such as Bacteroidetes on wild plants and Proteobacteria on domesticated plants (Perez-Jaramillo *et al*., 2018). The tomato EC and RT had increased Proteobacteria abundance while showing a decrease in Actinobacteria when compared to soils and ruderal plants. The loss of Actinobacteria abundance in tomato, a domesticated crop, compared to the soils and ruderal plants suggests that it could be a domestication trade-off, as previously suggested by a correlation between microbiome structure and host evolutionary history (Bouffaud *et al*. 2014; Peiffer *et al*. 2013; Redford *et al*. 2010). A comparison of maize, its ancestor teosinte, and other Poaceae rhizospheric microbiomes showed correlations between microbiomes and host evolutionary distances (Bouffad *et al*., 2014). The closer community distance of ruderal plants to their soils, when compared to tomatoes (Fig. 3), showed the tomato host genotype microbiome selection having a larger effect than soil, also lowering its overall α-diversity in a probable outcome of domestication trade-offs.

The tomato enriched rhizospheric bacteria, such as *Caulobacter, Rhizobium, Asticcacaulis*, *Sphingobium, Sphingomonas,* and *Novosphingobium,* are all Proteobacteria (Fig. S5A). These bacterial genera have been isolated from sources such as freshwater (Chen *et al*., 2013), soil (Costa *et al*., 2006), and rhizospheres (Young *et al*., 2008; Schreiter *et al*., 2014; Yang *et al*., 2016). The genera *Caulobacter* and *Asticcacaulis* are characterized by having at least one appendage or prostheca that protrudes from the cell envelope and can play a role in adhesion to solid substrates, a helpful attribute for the colonization of plant roots (Poindexter, 1981).

The presence of OTUs assigned to the families Sphingomonadaceae and Bradyrhizobiaceae in roots of *S. lycopersicum* has been reported previously (Larousse *et al*. 2017). Both Sphingomonadaceae and Bradyrhizobiaceae OTUs were reduced with plants inoculated with the pathogen *Phytophthora parasitica,* compared to healthy plants. We found an overrepresentation of some *Sphingobium* and *Rhizobium* species, suggesting that their abundance could be used as a plant health proxy since we did not observe root rot symptoms in any of our individuals (Satour & Butler 1967). Moreover, in a previous work describing tomato roots microbiomes, *Sphingomonas* and *Sphingobium* were detected in more than 50% of the 16S rRNA gene OTUs (Lee *et al*. 2016). *Sphingobium* has been observed as the dominant genus in tomato roots elsewhere (Pii *et al*. 2016).

Metagenomic profiling of the microbial communities showed that ruderal plants and soils have a similar composition of predicted proteins (Fig. 4C), differentiating them from tomato rhizospheres and highlighting the host-dependent selection. The enrichment of Proteobacteria in tomato is in line with enriched genes such as for motility and chemotaxis, widely distributed amongst α, β, and γ-Proteobacteria (Liu and Ochman, 2007). Motility traits are important for host colonization; this has been tested by mutagenesis in *Pseudomonas fluorescens* WCS36, reducing colonization efficiency of plant roots (de Weert *et al.,* 2002).

Other enriched proteins were diguanylate cyclase and CpaE, involved in biofilm formation and pili production in *Caulobacter crescentus* (Skerker & Shapiro, 2000), enriched in *S. lycopersicum* roots. Another interesting metabolic feature relevant for the plant-associated niche found in tomato roots is the enzyme xyloglucanase, which is involved in the degradation of xyloglucan, a heteropolysaccharide that comprises up to one-quarter of the total carbohydrate content of terrestrial plant cell walls (Scheller & Ulvskov, 2010).

Within the tomato core, we found multiple strategies to cope with nitrogen scarcity, such as cyanophycin biosynthesis genes. Cyanophycin is a reserve polymer (arginine and aspartate) regulating N and C and mediates N storage, providing bacterial fitness advantages under nitrogen fluctuations (Watzer & Forchhammer, 2018). Within the tomato core metagenome, allantoin degradation genes were found, which could be used as the sole N source to produce ammonia (Cruz-Ramos *et al*., 1997; Ma *et al*., 2016). Patatin-like proteins were also found in the tomato core metagenome; they are phospholipases originally described in potato, but with abundant homologs in bacteria (Banerji & Flieger, 2004). Bacteria use patatins to target host cell membrane as effectors via the type III secretion system (Finck-Barbançon *et al*., 1997; Sato *et al*., 2003; Phillips *et al*., 2003) and are activated by ubiquitin (Anderson *et al*., 2015). The eukaryotic patatins are known to have antimicrobial activities (*e.g., Phytophthora infestans* inhibition) (Bártová *et al*., 2019). Tomato and potato, both belonging to the family Solanaceae, interact with microbes via patatin and patatin-like proteins, and we will further explore plant-microbe interactions mediated by these proteins. In the core RT metagenome, we also found leucine aminopeptidases. Interestingly, leucine aminopeptidase A (LapA) is expressed in tomato after wounding and prevents foraging (*e.g., Manduca sexta* foraging tomato) (Fowler *et al*., 2009). LapA is also transcriptional and protein-responsive to microbial pathogens (Pautot *et al*., 1993; Pautot *et al*., 2001). The bacterial leucine aminopeptidases found in tomato metagenomes could be expanding the plant defensive response through LapA, but this is yet to be explored.

Domesticated plants such as tomato fit the two-step model for microbiome acquisition (Bulgarelli *et al.,* 2013). In contrast, ruderal plants had a larger taxa diversity than the source soil and a higher protein richness than tomatoes, although they were planted in the same soils. Plants have been domesticated since the Neolithic age some 10,000 years ago (Purugganan *et al.,* 2009), and genomic changes in microbes linked to domestication processes have been documented (genome reduction, insertion sequences, and transposition expansions), such as the enriched genes found in RT (Mira *et al.,* 2006). We hypothesize that domestication decreased the microbial diversity of the tomato root microbiome compared with that of grasses growing in the same soils. Plant domestication is targeted at meeting the requirements of humans, thereby decreasing plant genetic variability and generating crops dependent on humans (Bulgarelli *et al*., 2013; Doebley *et al*., 2006). Interestingly, the two-step model for root microbiota resembles the effects of reductive gene diversity in crop domestication (Doebley *et al*. 2006). Current agricultural management includes practices such as fertilizer-driven production, which decrease the importance of plant-microbe interactions when scavenging for nutrients (van der Heijden *et al*., 2008). The larger microbial diversity observed in ruderal plants shows the commitment of wild plants to their microbes, fostering plant-microbe relationships which are not observed in domesticated cultivars (Wissuwa *et al*., 2009). We have previously tested other non-domesticated plants, such as the aquatic carnivorous bladderwort *Utricularia gibba* (Alcaraz *et al*. 2016) and the bryophyte species *Marchantia polymorpha* and *M. paleacea* (Alcaraz *et al*. 2018); both showed less diversity in their root analogs (bladders, and rhizoids) than their soil sources, supporting the two-step model. The *Marchantia* microbiomes even allowed us to perform an extreme microbial selection due to the *in vitro* propagation of these plants, highlighting a reduced core of closely related microbial inhabitants (Alcaraz *et al*. 2018). Testing multiple plants, wild and domesticated, could reduce the gaps in understanding the microbiome structure loss as a domestication trade-off. Interestingly, the two-step model is not as descriptive with metagenome-predicted proteins, and it probably needs further refinement maybe through linking the OTU abundance with pan-genomics and metagenomics to describe the genomic coding diversity (Delmont & Eren, 2018).

Describing the tomato core microbiome and metagenome under multiple soils also allowed us to test the plant genotype filtering effect, evaluating selected microbes in diverse environments. With the current advances in synthetic biology, the tomato core metagenome could lead to a tomato root metagenomic “chassis”. This, in turn, could lead to microbe-complemented plant breeding programs aiming to reduce and optimize fertilizer use while increasing plant resilience such as that observed in ruderal plants. Further possibilities could be the recovery of the domesticated missed root microbes from wild plants.

### Concluding remarks

By using 16 geochemically diverse soils as microbial inputs for root colonization, we discarded the role of soil as the major structuring factor of root microbial communities, particularly of their coding genes. Further work is needed for detecting other environmental microbe sources than the soil for rhizosphere metagenomic diversity. Weather-dependent ruderal plant roots are a nutrient and moisture oasis for soil microbial communities with a higher taxonomic α-diversity. The tomato root microbiome followed the two-step model of microbiome acquisition. The reduced total protein number, along with significant enrichments in the tomato root metagenomes compared to ruderals and soils, suggests a tomato rhizosphere specialization and a possible domestication trade-off. Our experimental setup showed that tomato enriches plant-microbe interaction genes. Altogether, our results show that tomato roots have a convergent, genotype driven, and reduced microbiome compared to their source soils, following the two-step selection model for the root microbiome. This is contrary to the ruderal plants, which exhibit a larger microbiome diversity than their soils, not following the two-step model.

## Methods

### Soil and local plant roots sampling

Edaphological charts were used to locate 8 different soil groups, according to the United Nations FAO classification (IUSS, 2015) from 16 different geographic locations described in Fig. 1A, and Table 1. In each location, 0.09 m^2^ quadrats were placed, and duplicate root samples were taken from the quadrat dominating plant species, along with the soil below them. We collected 2 kg of each soil into sterile plastic bags for the common garden experiment and biogeochemical analysis. All soil samples were taken from a depth not larger than 10 cm. Soil was kept at 4°C in a darkroom until greenhouse experiments were conducted. *In situ* soils were collected, for each soil group, and poured into duplicate sterile centrifuge tubes (50 mL volume), then immediately field frozen in liquid nitrogen until storage into a −80°C freezer, until metagenomic DNA extraction.

### Common garden experiment, harvesting, and sample collection

The tomato seeds used were *Solanum lycopersicum* L. Cv. *Río grande* (Sun Seeds, Parma, ID, USA). Seeds were surface disinfected in 70% ethanol for 1 min, followed by a wash in 2.5% NaOCl for 2 min, and rinsed with sterile distilled water. Seeds were germinated in 1% agar for 96 h in a dark growth chamber at 27°C. Sprouts were aseptically transplanted into duplicated pots filled with the collected soils, two plants per pot were transplanted, summing 4 biological replicates; additionally, pots with each soil were prepared without plants (US, Fig. 1C). Pots were set in the greenhouse randomly, and plants were watered with tap water every other day and harvested after 60 days of growth. All soil samples (Fig. 1) were collected in 50 mL sterile tubes and frozen at −80°C until metagenomic DNA extraction.

Roots were separated from shoots to collect rhizosphere and endosphere samples by removing loose soil, followed by a washing and ultrasound procedure in 1X PBS buffer (137 mM NaCl; 2.7 mM KCl; 10 mM Na_2_HPO_4_; 1.8 mM KH_2_PO_4_) as described before (Lundberg *et al*., 2012). Tomato rhizosphere and endosphere metagenomic pellets, were recovered through centrifugation (50mL tubes centrifuged at 1,300 g during10 min). Roots and shoots were oven-dried at 60°C for 24 hours to measure plant biomass production. Due to low DNA extraction efficiency by this method in ruderal plant roots, they were cut and separated into ten 1.5mL tubes, which received the same treatment as the 50 mL tubes. All sample pellets were frozen and kept −80°C until metagenomic DNA extraction (Fig 1).

### Soil geochemical analyses

Initial and final soils were oven-dried for 24 h at 70°C. The pH was measured in deionized water (1:4 w:v) with a Corning digital pH meter. Total carbon was measured by coulometric combustion detection (Huffman 1977) with a Total Carbon Analyzer (UIC Mod. CM 5012; Chicago, IL, USA), total nitrogen was determined by a semi-Kjeldahl method and phosphorus by the molybdate colorimetric method after ascorbic acid reduction (Murphy and Riley, 1962) using a Bran-Luebbe Auto Analyzer III (Norderstedt, Germany). The Lang’s aridity index (Lang, 1920) of each site was calculated using historical data of mean annual precipitation and temperature for each sampling location, and data was consulted at Atmospheric Sciences Center of UNAM (http://uniatmos.atmosfera.unam.mx/ACDM/). Nonmetric multidimensional scaling (NMDS) of the samples was calculated with the geochemical data using the metaMDS function in the vegan R package (Oksanen *et al*., 2015) and plotted with ggplot2 (Wickham, 2009). Detailed statistical and bioinformatic methods are available at https://github.com/genomica-fciencias-unam/Barajas-2020.

### Metagenomic DNA processing and massive sequencing

The metagenomic DNA of all samples was extracted using the Mobio PowerSoil DNA extraction kit (MoBio, Carlsbad, CA, USA), following the manufacturer’s instructions. Briefly, for soils, approximately 0.25g were used for the extraction, for rhizosphere and endosphere pellets collected after washing and sonication of the roots were used respectively, as previously described (Lundberg *et al.,* 2012). Then, the Mobio protocol was slightly modified to get extra DNA by heating the C6 elution solution to 60°C before eluting the DNA, and two 30 uL elutions were performed on the same spin filter. The same DNA was used for both amplicon and whole metagenome shotgun sequencing.

PCR amplification of the 16S rRNA gene was performed in duplicates, followed by the Illumina® MiSeq protocol for 16S metagenomic sequencing library preparation (Illumina 2013) using the 341F/805R primer pair targeting the V3-V4 regions with the Illumina sequencing adaptors in 5’ (341F: 5’-CCTACGGGNGGCWGCAG*-*3’; 805R: 5’- ACTACHVGGGTATCTAATCC 3’). PCR reactions were performed in a 20 µL volume, consisting of 0.16 µL *Pfx* polymerase (0.02U/µL) (Invitrogen, Thermo Fisher Scientific, Waltham, MA) 2µL buffer, 3 µL enhancer, 1.2 µL of each primer (5µM), 1.6 µL dNTPs (2.5mM), 0.6 µL Mg2S04 (1.5µM), 9.2 µL PCR grade water and 2 µL DNA template. The PCR program for amplification was 95°C for 3 min, followed by 5 cycles of 94°C for 30 s, 55°C for 30 s, 68°C for the 30s, followed by 25 cycles of 94°C for 5 s and 68°C for 30 s. The duplicate amplification products of each sample were pooled and purified with the SV Wizard PCR Purification kit (Promega, Madison, WI) following the manufacturer’s instructions. Amplicon library sequencing was done in the Illumina® MiSeq platform in a 2×300 paired-end configuration at the University Unit of Massive Sequencing and Bioinformatics (http://www.uusmd.unam.mx) of the Biotechnology Institute, UNAM, Mexico. Whole shotgun metagenome sequencing libraries were prepared using the Truseq PCR free library preparation kit for selected initial soils, ruderal plants, and *S. lycopersicum* rhizospheres, which were then sequenced with an Illumina HiSeq 2000 in a 2 x 100 bp reads, at the facilities of Macrogen, Korea (https://www.macrogen.com).

### 16S rRNA gene amplicon sequence analysis

The 16S rRNA protocol used in this work had been used previously and is detailed at GitHub (Alcaraz *et al.,* 2018; https://genomica-fciencias-unam.github.io/SOP/). In summary, gene amplicon libraries were quality inspected using Fastx Toolkit (http://hannonlab.cshl.edu/fastx_toolkit/) and trimmed to a 250 bp length. Trimmed paired-end reads were assembled using Pandaseq (Masella *et al*., 2012). The assembly was performed using a minimum overlap of 15 bp, the minimum output length of 250 bp, the maximum output length of 470 bp, and an alignment threshold of 95%. Finally, assembled sequences were filtered using a minimum PHRED score of 20. All the samples were concatenated and clustered into OTUs, using a 97% identity threshold with *cd-hit-est* (Li *et al*., 2006). The taxonomy of representative sequences was assigned against Greengenes (De Santis *et al*.,2006) database with QIIME’s scripts (Caporaso *et al*. 2010). After taxonomic classification, singletons, and chimeras were removed as well as sequences corresponding to the mitochondria, chloroplast, and unassigned hits were filtered out. Finally, the representative OTU sequences were aligned with SSU-align (Nawrocki, 2009), and a phylogenetic tree was constructed with Fasttree (Price *et al*., 2009). Detailed statistical and bioinformatic methods are available at https://github.com/genomica-fciencias-unam/Barajas-2020.

### Metagenomic shotgun sequence analysis

The quality control of whole shotgun metagenome sequences was done using Trimmomatic (Bolger *et al*., 2014), only paired-end matched reads were used for subsequent analysis. We filtered out metagenomic reads matching *S. lycopersicum* genome, while soils and ruderal plants rhizosphere libraries were filtered against the *Oryza sativa* genome with Bowtie2 (Langmead and Salzberg, 2012). Quality and host filtered metagenomic libraries were used to assemble individual metagenomes with metaSPADES (Nurk *et al*., 2017). High-quality reads were mapped against the metaSPADES contigs, and unmapped reads were subjected to a second assembly with Velvet (Zerbino *et al*., 2008). The resulting contigs from both assemblies were merged and used to predict ORFs and coding proteins with Prodigal (Hyatt *et al*., 2010). Annotation of predicted proteins was made against the M5NR database (Wilke *et al*., 2012) using DIAMOND (Buchfink *et al*., 2015) with the following parameters -f6 -e 1e-10 -k 10 -p1, retrieving Refseq (Pruitt *et al*., 2007) and SEED subsystems (Overbeek *et al*. 2014) annotations from M5NR matched identifiers. The abundance of each predicted protein was calculated by mapping the high-quality reads against the predicted ORFs with Bowtie2. All the predicted proteins were clustered using cd-hit (Li *et al*., 2006) using a 70% identity threshold, and then parsed into a biom formatted matrix, used as input for sets comparison using UpSetR (Conway and Gehlenborg, 2017). The binning of whole shotgun metagenomic reads was performed with Kaiju (Menzel *et al*., 2016). Detailed statistical and bioinformatic methods are available at https://github.com/genomica-fciencias-unam/Barajas-2020.

### Diversity analysis

The α and β-diversity of soils, rhizospheres, and endospheres from each site were calculated with phyloseq (McMurdie & Holmes, 2013), and vegan R (Oksanen *et al*., 2015) packages. Taxonomic β-diversity was assessed using a weighted Unifrac (Lozupone *et al*., 2006) distance matrix. Then, microbiomes were hierarchically clustered with the *hclust* method using complete distances and clustering evaluated through the ANOSIM function. OTUs were clustered at the genus level, and Venn diagrams were used to compare the complete root system (rhizosphere + endosphere) microbiome composition of ruderal plants, *S. lycopersicum,* and initial soils using the web interface https://bioinformatics.psb.ugent.be/webtools/Venn/. Unique soil, ruderal plants, *S. lycopersicum,* and the ruderal plants-*S. lycopersicum* intersection taxonomic profiles were described at the phylum level based on OTU abundances.

Metabolic β-diversity was estimated through a constrained analysis of principal coordinates (CAP) analysis using Bray-Curtis dissimilarity based on the total abundance of predicted proteins. Differential OTUs and protein abundances comparing rhizospheres or endosphere against soils were calculated using DESeq2 (Love *et al*., 2014) with a Wald statistical test and a local fit of the data. For 16S rRNA data, OTUs were considered differentially abundant between groups using a *p* < 0.01, for metagenome predicted proteins, a *p* < 0.001 was used as a cut-off. The collected ruderal plant species were identified by their 16S rRNA matches to NCBI’s NR database representing a variety of 5 different plant families, mainly grasses (Poaceae N=10, Asteraceae N=3, Lamiaceae N=1, Fabaceae N=1, and Fagales N=1; Fig. S1). Detailed statistical and bioinformatic methods are available at https://github.com/genomica-fciencias-unam/Barajas-2020.

## Acknowledgments

We want to thank Rodrigo Velázquez Durán for his technical support with the soil physicochemical properties characterization. Selene Molina and Jazmín Blaz for their help in sample processing, and Francisco González, who helped in the sampling of soils, and ruderal plants’ roots. We also thank Rodrigo García Herrera at LANCIS-Instituto de Ecología, UNAM, for his support in HPC computing.

## Author Contributions

HB, LS, MP, RC, FG, and LA conceived and designed the research, HB and LA wrote the paper, HB, SM, MP, and LA performed field sampling, HB and SM performed experiments, HB, SM, CH, MR, LA analyzed data, MP, RC, FG, and LA provide reagents and material, HB, MP, and LA created figures, all the authors read and approved the manuscript.

## Data availability

Raw sequence reads for the whole study were deposited in the NCBI. The BioProject database under the accession codes PRJNA603586 for soils, and PRJNA603590 for rhizosphere, and endosphere 16S rRNA gene libraries. Whole shotgun metagenomes can be found under BioProject accession code PRJNA603603.

## Supplementary material

Complete supplemental material available at: https://github.com/genomica-fciencias-unam/Barajas-2020

Complete supplemental material available at: https://github.com/genomica-fciencias-unam/Barajas-2020

## Supplementary Figures

**Figure S1.**
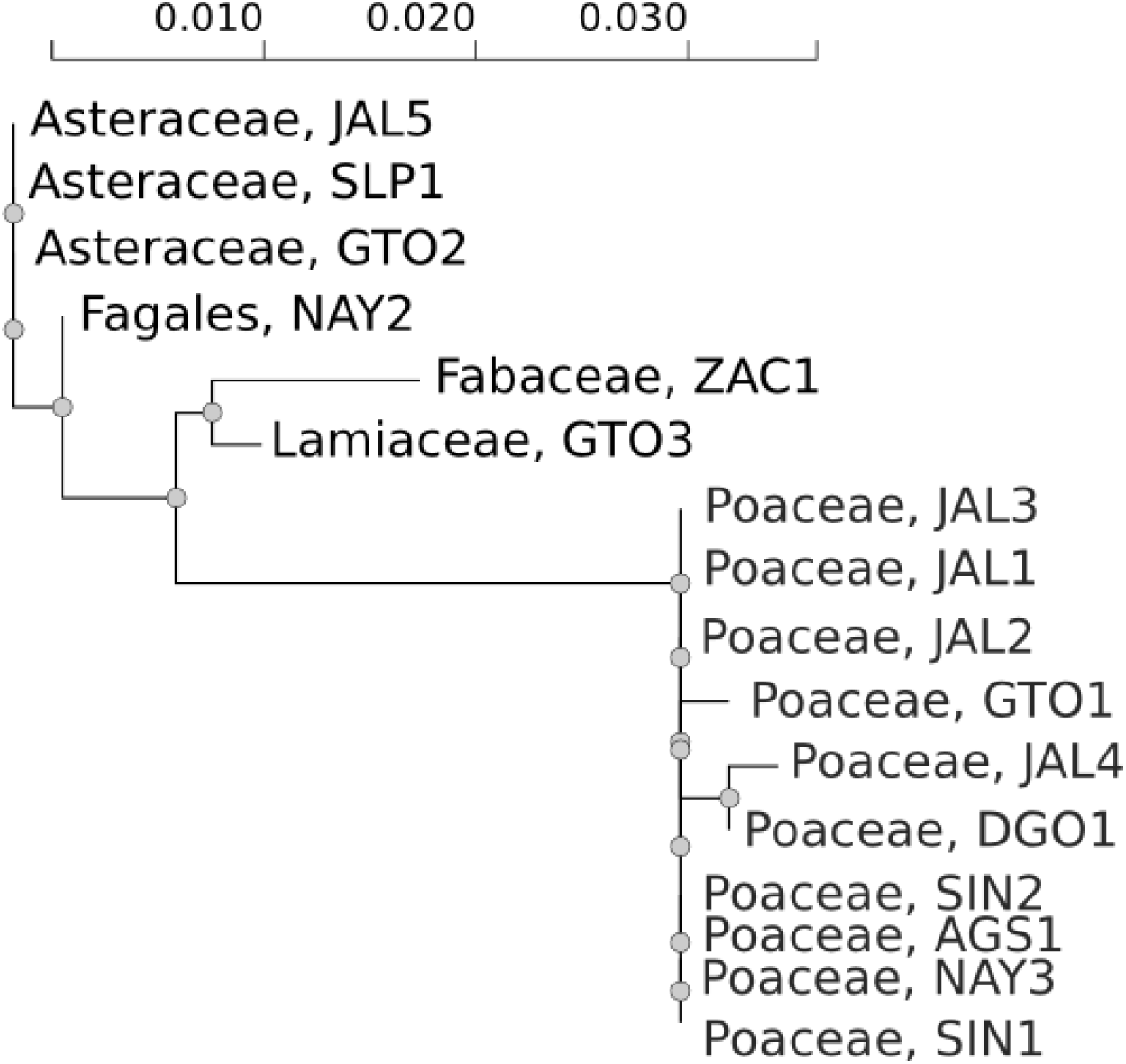
Taxonomic identification of the ruderal plants using the 16S rRNA gene. The maximum likelihood phylogeny and best hit classification showed that the ruderal plants collected in this work, were mainly grasses (*Poaceae*, N=11), composite (*Asteraceae*, N=3), and then single representatives of *Fagales*, *Fabaceae*, and *Lamiaceae*.

**Figure S2.**
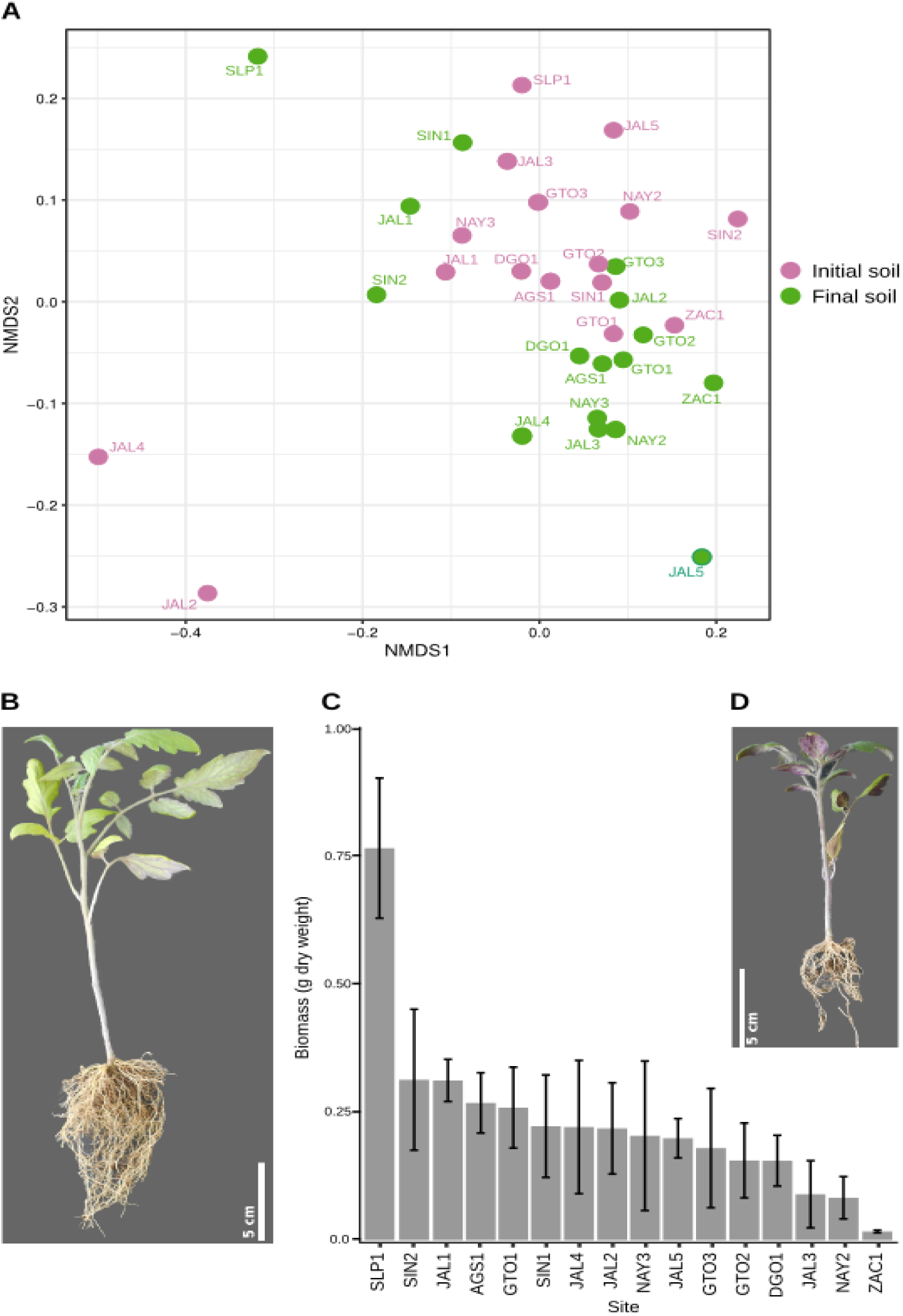
Geochemical diversity of source and final soil, and plant biomass production in common garden experiment. **A)** NMDS ordination bi-plot of initial soils (SI) and final soils (FS) calculated with the following soil abiotic properties: Aridity index, total carbon content, total nitrogen content, total phosphorus content. NMDS stress=0.116. **B)** Example of tomato individual grown on SLP1 soil. **C)** Bar-plot showing average biomass production of tomato plants in the common garden experiment. **D)** Example of tomato individual grown on NAY2 soil.

**Figure S3.**
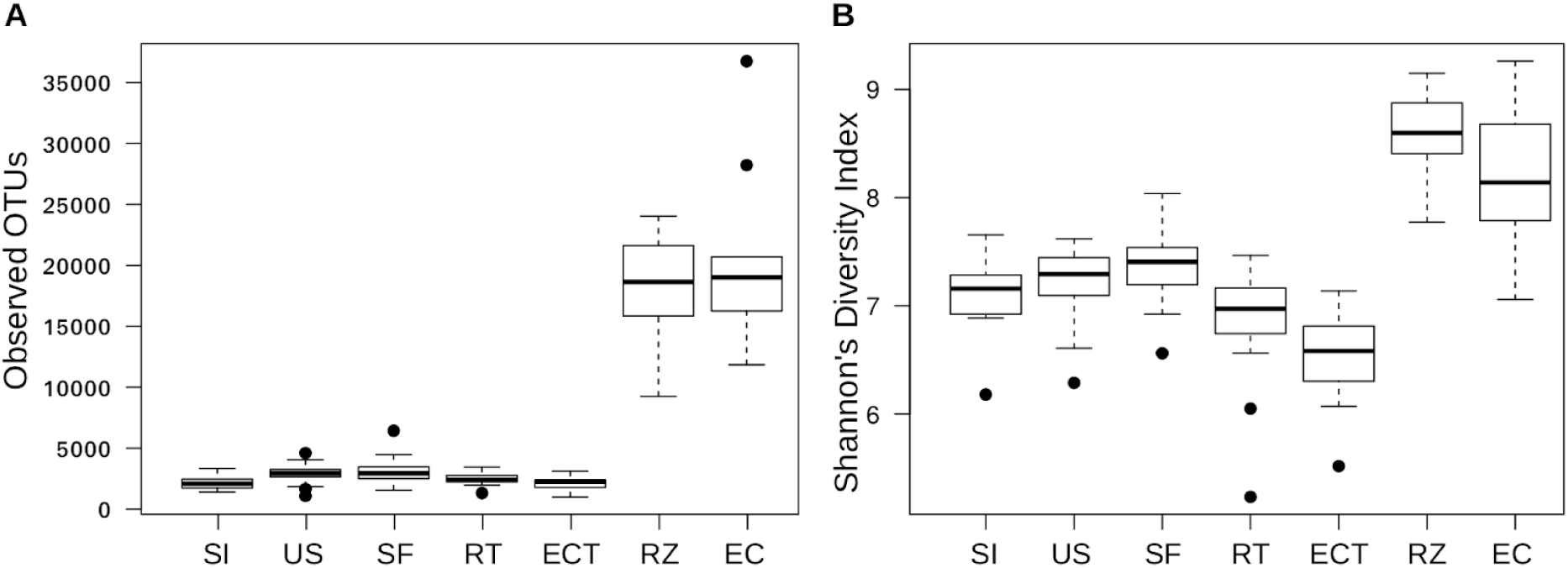
Richness and diversity of initial, final, and unplanted soil, tomato, and ruderal plants rhizosphere and endosphere. Boxplots showing median values of **A)** Observed OTUs and **B)** Shannon diversity index. SI=initial soil, FS=Final soil, US=unplanted soil, RT=tomato rhizosphere, ECT=tomato endosphere, RZ=ruderal plants rhizosphere, EC=ruderal plants endosphere.

**Figure S4.**
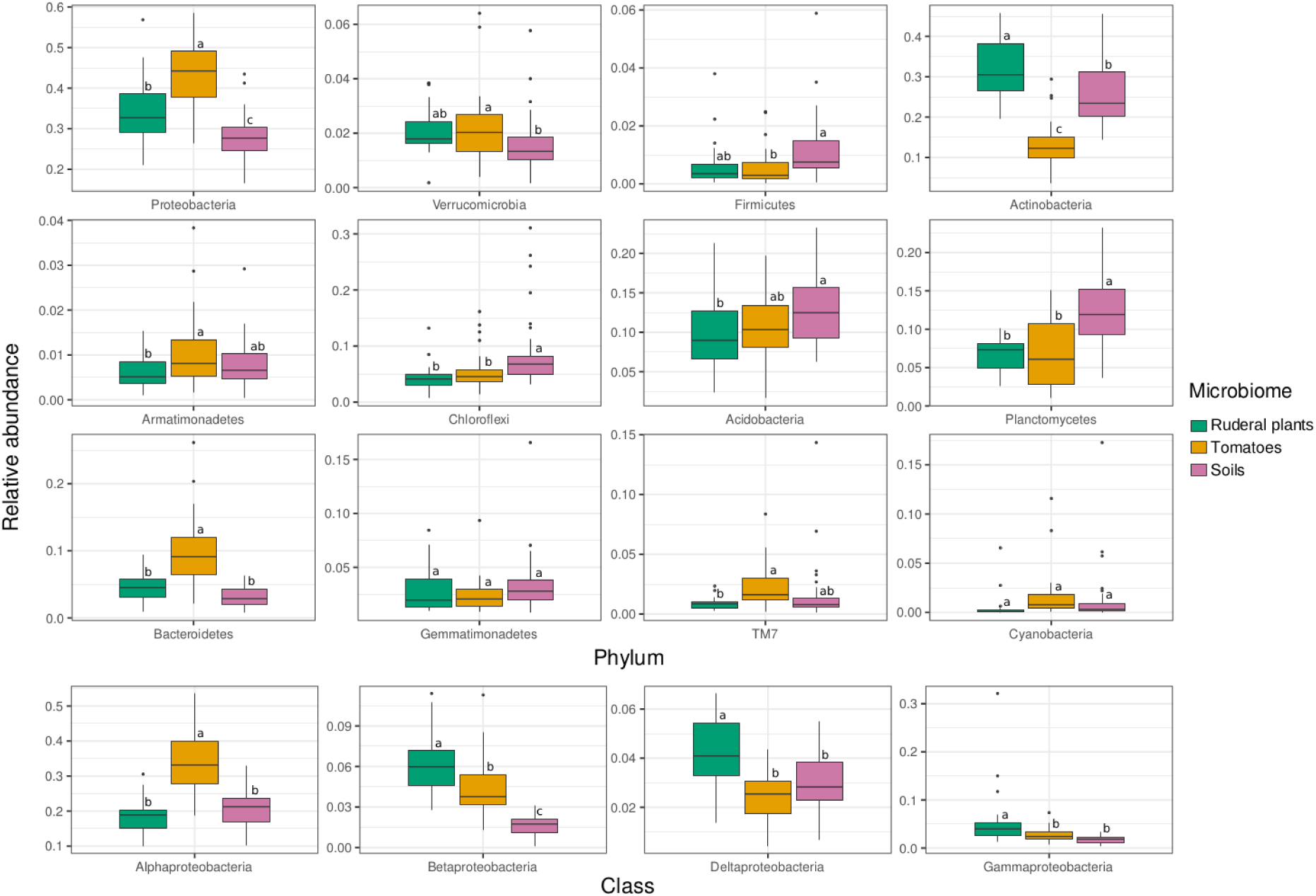
Relative abundance of bacterial phyla in soils, ruderal plants and tomato roots. Each panel shows the relative abundance of different phyla. Proteobacteria are shown at the class level.

**Figure S5.**
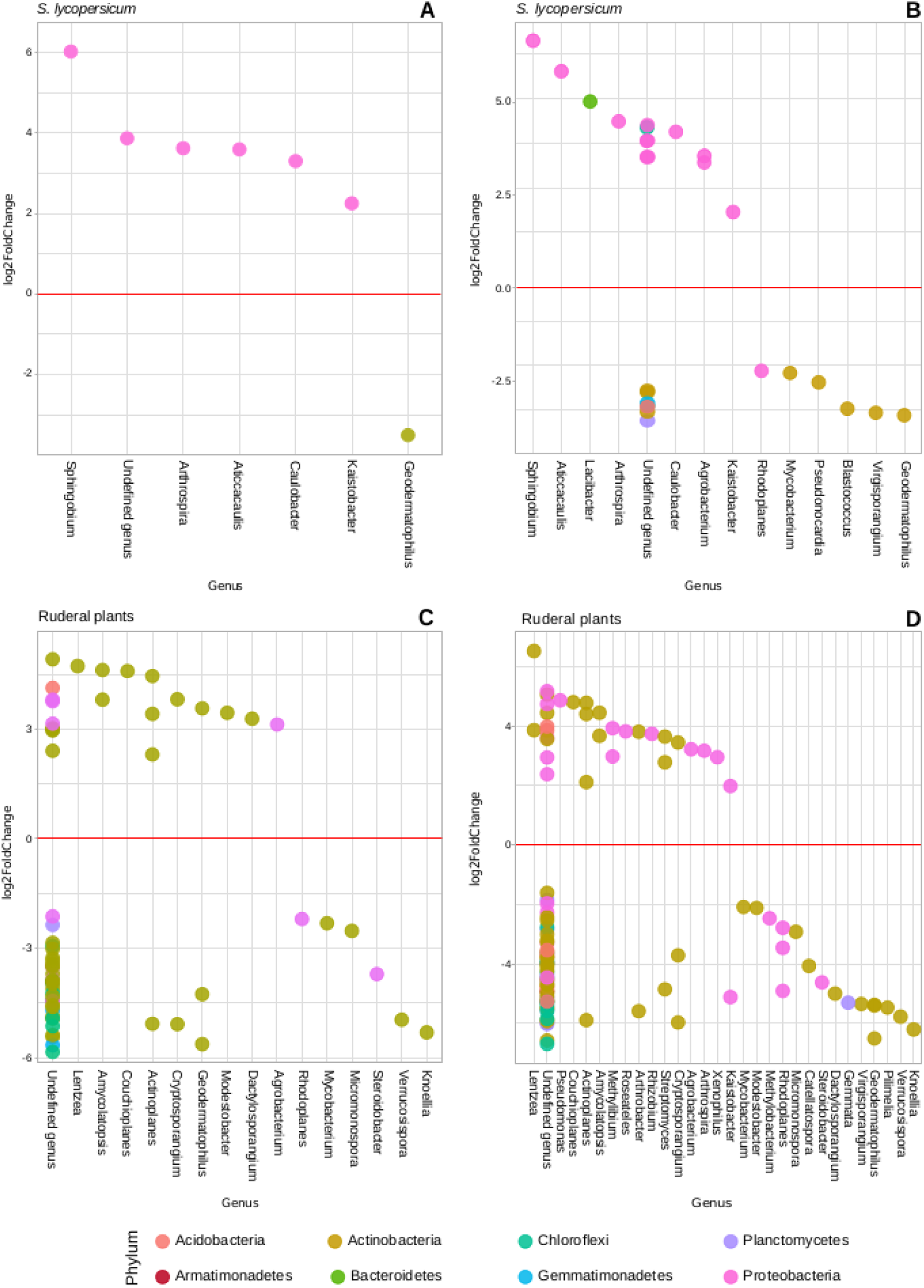
Enriched OTUs in *S. lycopersicum* and ruderal plants root systems. Log2 Fold Change of OTUS abundance in comparisons between initial soils and **A)** Tomato rhizospheres. **B)** Tomato endosphere. **C)** Ruderal plants rhizosphere. **D)** Ruderal plants endosphere. Positive Log2 Fold Change values indicate differentially abundant OTUs in plant roots systems.

**Figure S6.**
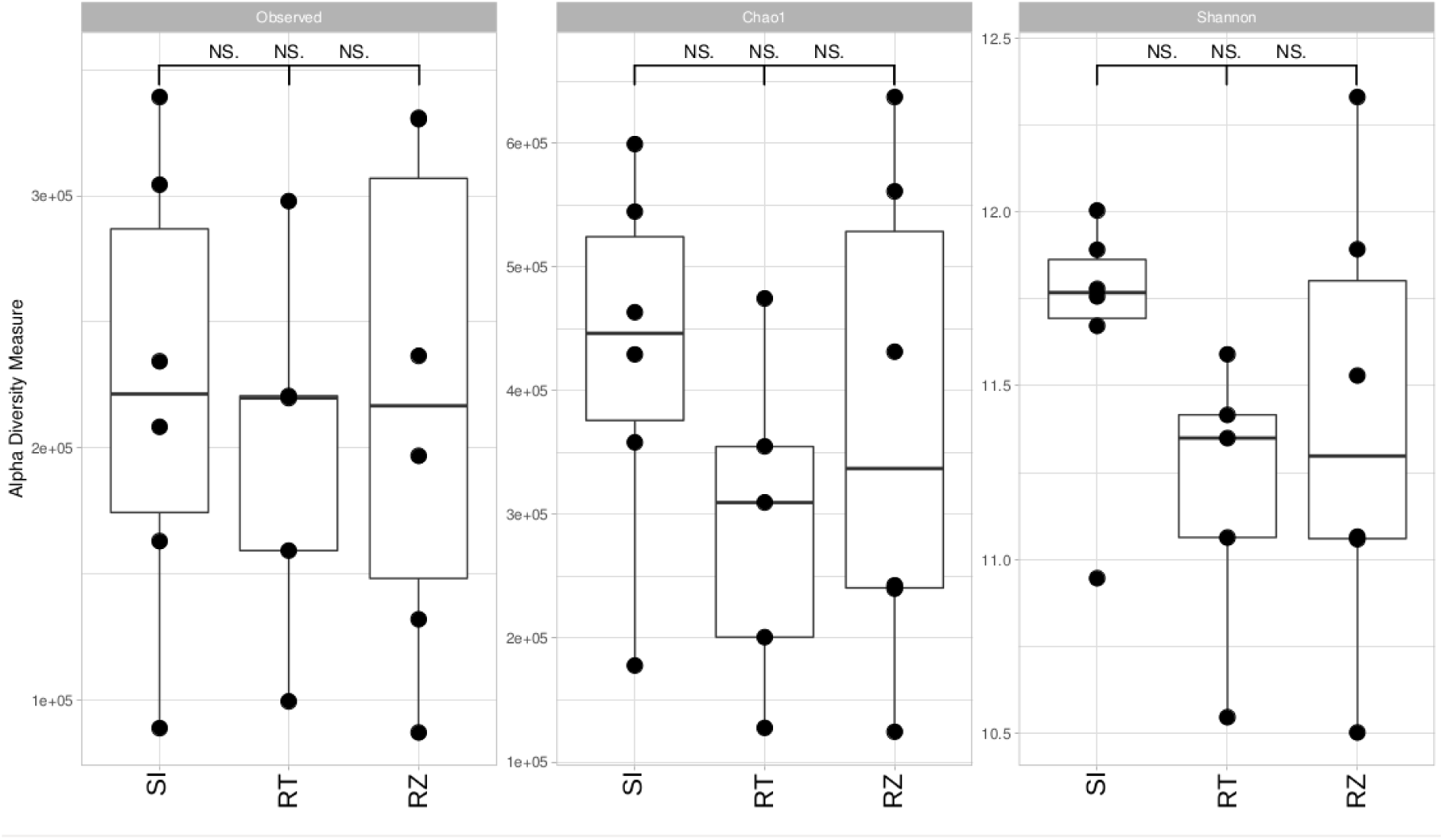
Alpha diversity of predicted proteins in soil sources, *S. lycopersicum*, and ruderal plants metagenomes. The number of observed proteins, Chao1, and Shannon diversity index are shown. No significant differences were found between groups.

**Figure S7.**
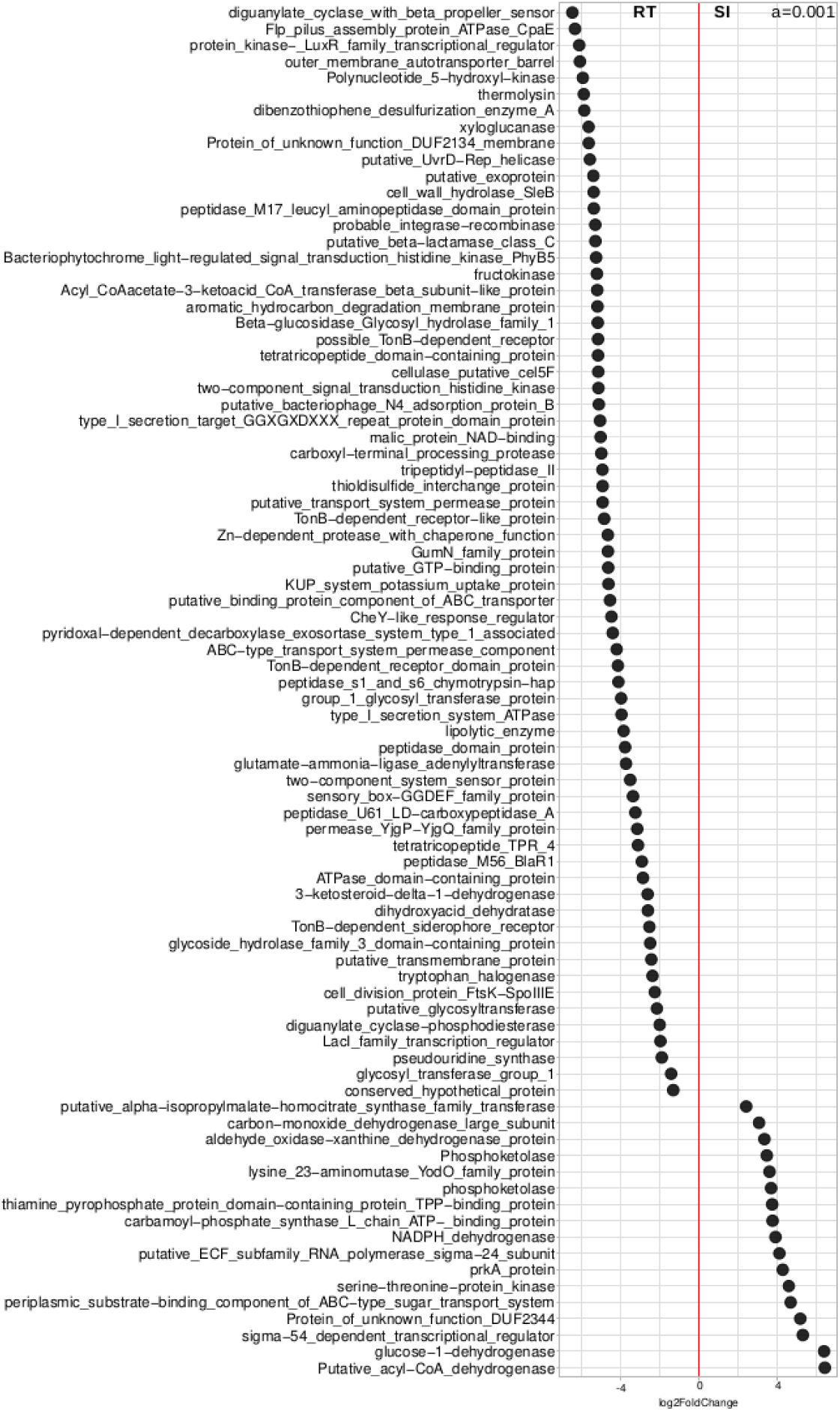
Differentially abundant proteins in *S. lycopersicum* rhizosphere against soil metagenomes. Log2 fold change values (*p* < 0.001) of annotated proteins are shown. Sixty-four proteins were differentially abundant in *S. lycopersicum* rhizosphere. Negative log2 fold change values are tomato rhizosphere enriched proteins, while positive values represent soil enriched proteins.

**Figure S8.**
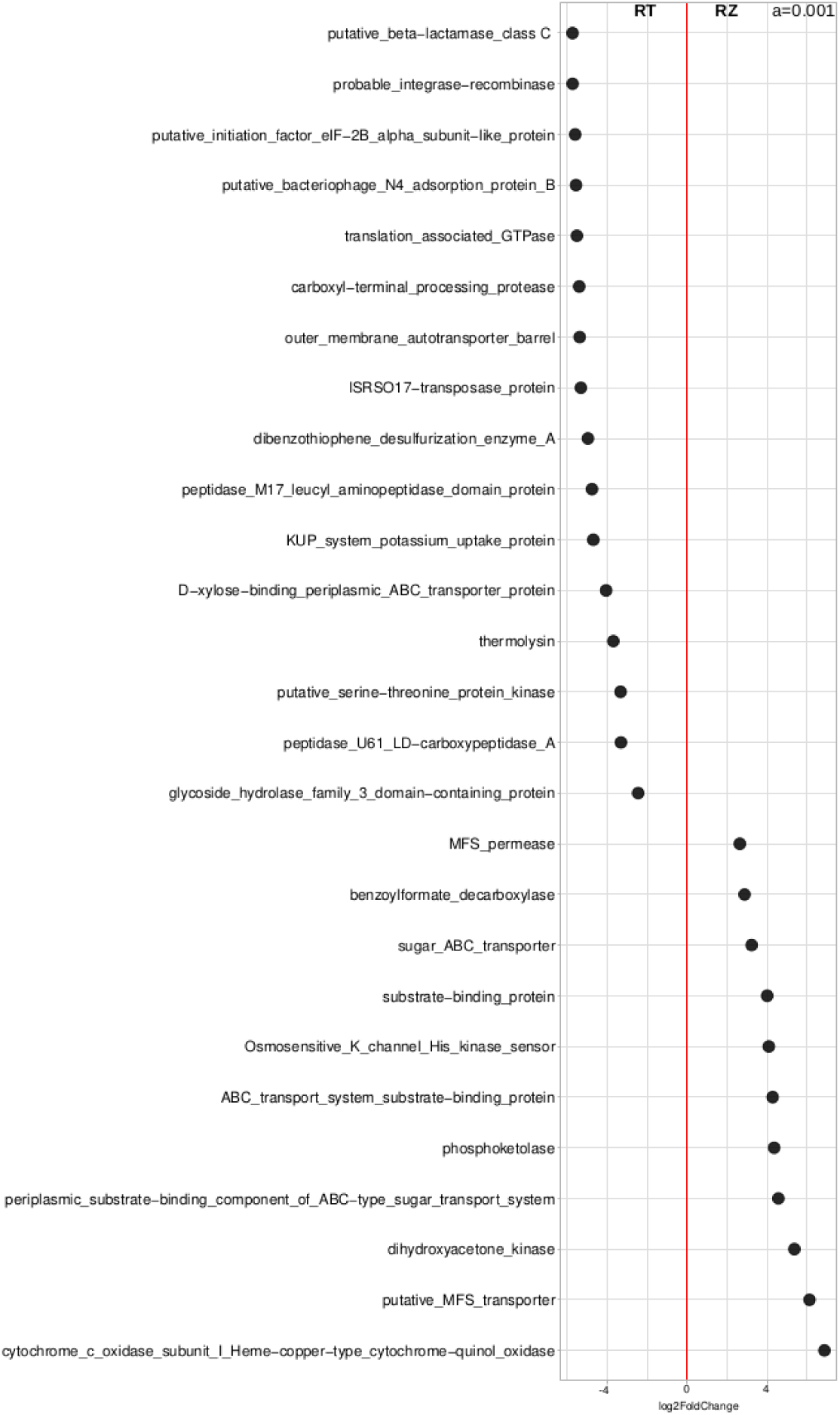
Differentially abundant proteins in the comparison between ruderal plants and *S. lycopersicum* rhizosphere metagenomes. Log2 fold change (*p* < 0.001) of annotated proteins are shown. Sixteen proteins were differentially abundant in tomatoes and eleven in ruderal plants rhizospheres. Negative log2 fold change values are tomato rhizosphere enriched proteins, while positive values are enrichments in ruderal plants.

**Figure S9.**
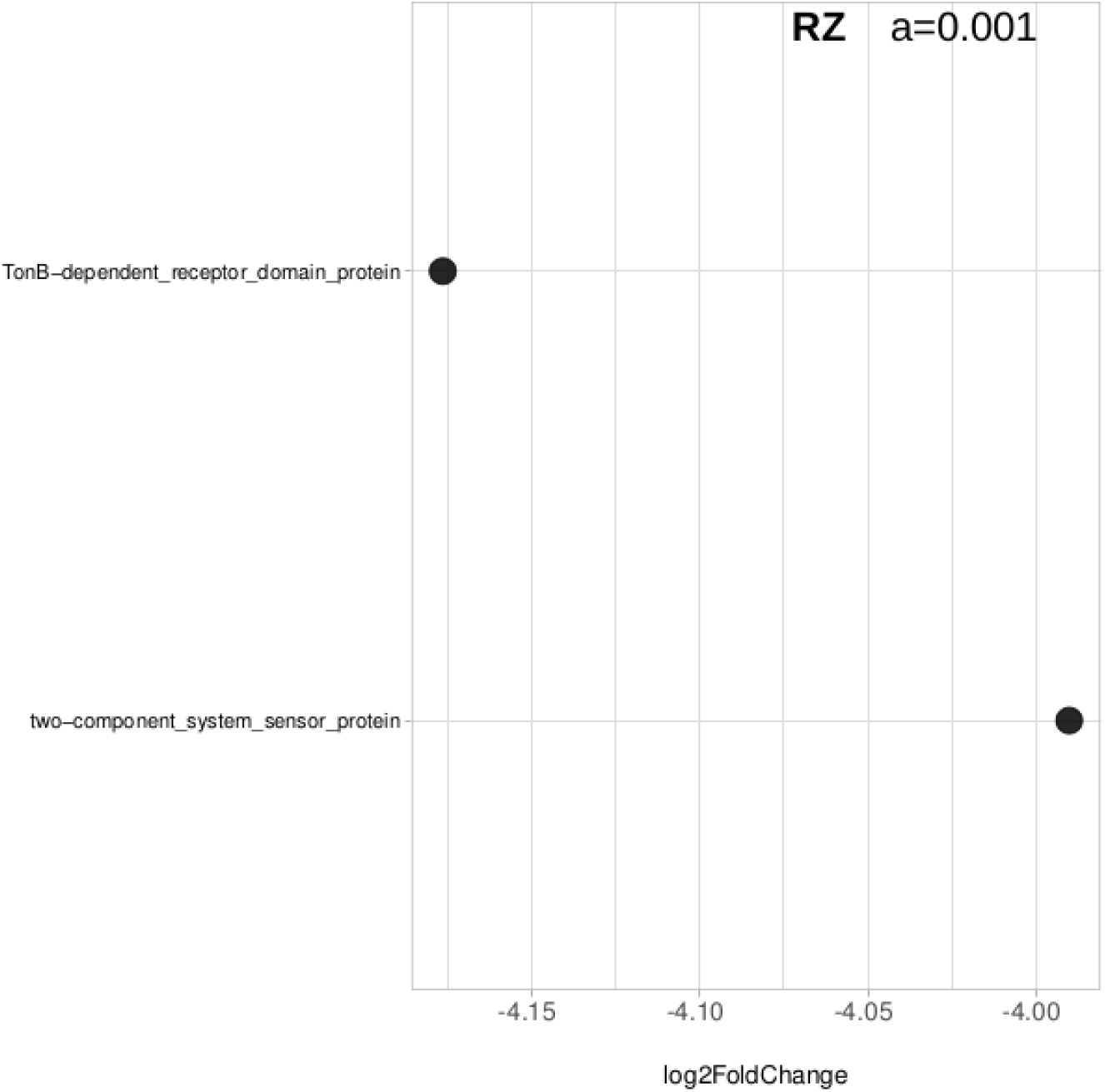
Differentially abundant proteins in ruderal plants rhizosphere against initial soil metagenomes. Log2 fold change (*p* < 0.001) of annotated proteins are shown. Only two proteins were differentially abundant in ruderal plants’ rhizosphere. Negative Log2 fold change values are proteins enriched in ruderal plants, positive are enriched in soil.

**Figure S10.**
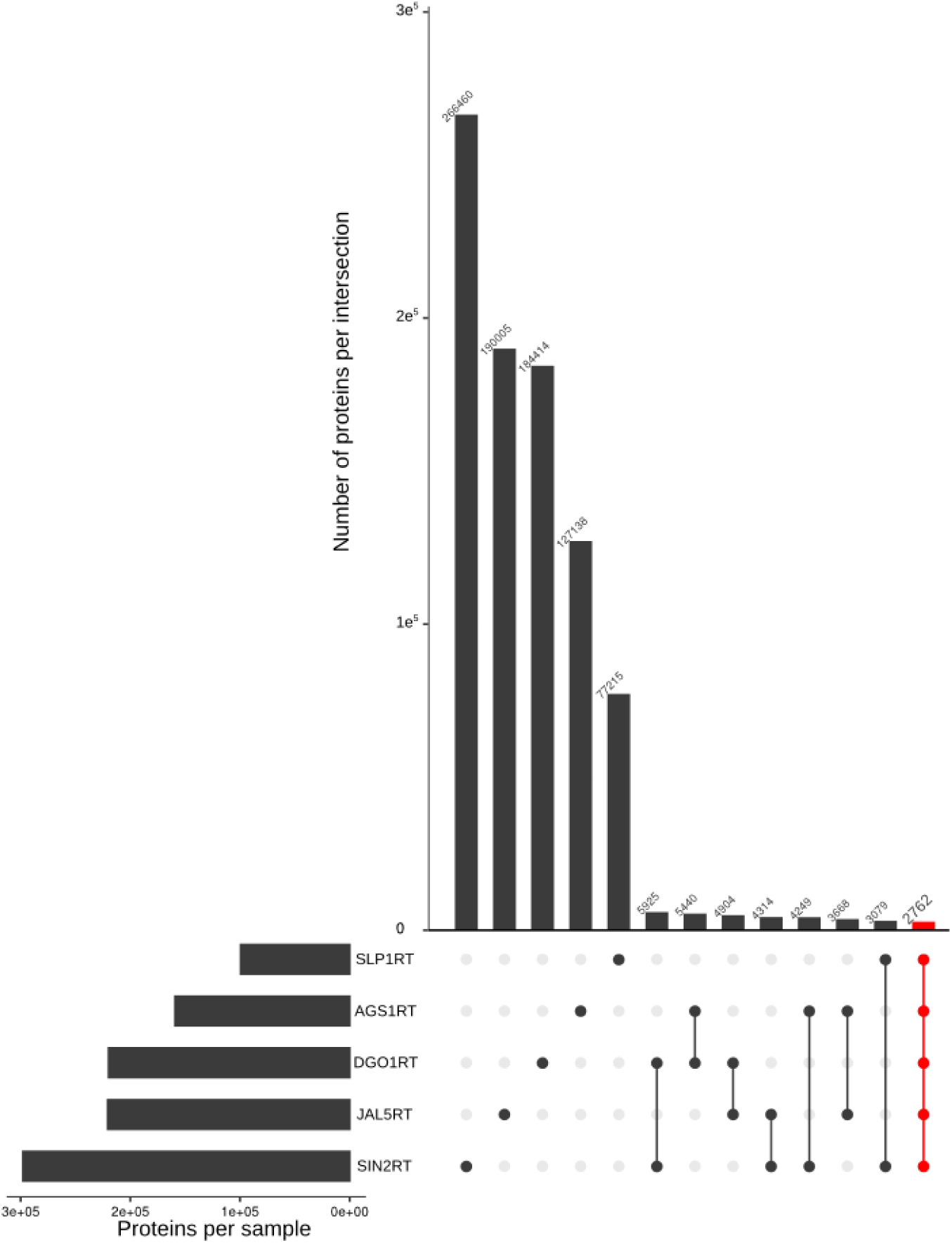
Tomato core metagenome. Upset diagram showing unique and shared sets of proteins in the tomato rhizosphere metagenomes. The tomato core metagenome consists of 2,762 proteins.

## Supplementary tables

**Supplementary Table S1.** Summary of paired-end reads and assembled sequences in 16S rRNA gene libraries.

**Supplementary Table S2.** Diversity metrics of microbiome samples.

**Supplementary Table S3.** Unique and shared OTUs between tomato, ruderal plants and soils.

**Supplementary Table S4.** Pairwise cophenetic distances between microbiomes.

**Supplementary Table S5.** Phylum relative abundance in soils, rhizospheres and endospheres of tomato and ruderal plants.

**Supplementary Table S6.** Deseq2 enriched OTUs in soils, rhizospheres and endospheres of tomato and ruderal plants.

**Supplementary Table S7.** Summary of whole shotgun metagenomes sequencing and assembly.

**Supplementary Table S8.** Shared and unique predicted proteins between tomato, ruderal plants and soils metagenomes.

**Supplementary Table S9.** Whole shotgun metagenomes α-diversity metrics.

**Supplementary Table S10.** Abundance of taxa based on binning of metagenomic reads with Kaiju.

**Supplementary Table S11.** Deseq2 enriched proteins in soils, rhizospheres and endospheres of tomato and ruderal plants.

**Supplementary Table S12.** Soils, tomato, and ruderal plants rhizospheres core metagenomes.

**Supplementary Table S13.** Shared and unique proteins between soil, tomato, and ruderal plants core metagenomes.

## References

1. Alcaraz, L. D., Martínez-Sánchez, S., Torres, I., Ibarra-Laclette, E. and Herrera-Estrella, L. (2016) ‘The metagenome of *Utricularia gibba*’s traps: Into the microbial input to a carnivorous plant’, PLoS ONE, 11(2), p. e0148979. doi: 10.1371/journal.pone.0148979.

2. Alcaraz, L. D., Peimbert, M., Barajas, H. R., Dorantes-Acosta, A. E., Bowman, J. L. and Arteaga-Vázquez, M. A. (2018) *‘Marchantia* liverworts as a proxy to plants’ basal microbiomes’, Scientific Reports, 8(1), p. 12712. doi: 10.1038/s41598-018-31168-0.

3. Anderson, D. M., Sato, H., Dirck, A. T., Feix, J. B. and Frank, D. W. (2015) ‘Ubiquitin activates patatin-like phospholipases from multiple bacterial species’, Journal of Bacteriology. Edited by G. A. O’Toole, 197(3), pp. 529–541. doi: 10.1128/JB.02402-14.

4. Bais, H. P., Weir, T. L., Perry, L. G., Gilroy, S. and Vivanco, J. M. (2006) ‘The role of root exudates in rhizosphere interactions with plants and other organisms’, Annual Review of Plant Biology, 57(1), pp. 233–266. doi: 10.1146/annurev.arplant.57.032905.105159.

5. Banerji, S. and Flieger, A. (2004) ‘Patatin-like proteins: a new family of lipolytic enzymes present in bacteria?’, Microbiology, 150(3), pp. 522–525. doi: 10.1099/mic.0.26957-0.

6. Bártová, V., Bárta, J. and Jarošová, M. (2019) ‘Antifungal and antimicrobial proteins and peptides of potato (*Solanum tuberosum* L.) tubers and their applications’, Applied Microbiology and Biotechnology, 103(14), pp. 5533–5547. doi: 10.1007/s00253-019-09887-9.

7. Baudoin, E., Benizri, E. and Guckert, A. (2002) ‘Impact of growth stage on the bacterial community structure along maize roots, as determined by metabolic and genetic fingerprinting’, Applied Soil Ecology, 19(2), pp. 135–145. doi: 10.1016/S0929-1393(01)00185-8.

8. Berendsen, R. L., Pieterse, C. M. J. and Bakker, P. A. H. M. (2012) ‘The rhizosphere microbiome and plant health’, Trends in Plant Science, 17(8), pp. 478–486. doi: 10.1016/j.tplants.2012.04.001.

9. Bolger, A. M., Lohse, M. and Usadel, B. (2014) ‘Trimmomatic: A flexible trimmer for Illumina sequence data’, Bioinformatics, 30(15), pp. 2114–2120. doi: 10.1093/bioinformatics/btu170.

10. Bouffaud, M. L., Poirier, M. A., Muller, D. and Moënne-Loccoz, Y. (2014) ‘Root microbiome relates to plant host evolution in maize and other *Poaceae**’*, Environmental Microbiology, 16(9), pp. 2804–2814. doi: 10.1111/1462-2920.12442.

11. Boyd, E. S. and Peters, J. W. (2013) ‘New insights into the evolutionary history of biological nitrogen fixation’, Frontiers in Microbiology, 4(AUG), pp. 1–12. doi: 10.3389/fmicb.2013.00201.

12. Brader, G., Compant, S., Mitter, B., Trognitz, F. and Sessitsch, A. (2014) ‘Metabolic potential of endophytic bacteria’, Current Opinion in Biotechnology, 27, pp. 30–37. doi: 10.1016/j.copbio.2013.09.012.

13. Buchfink, B., Xie, C. and Huson, D. H. (2015) ‘Fast and sensitive protein alignment using DIAMOND’, Nature Methods, 12(1), pp. 59–60. doi: 10.1038/nmeth.3176.

14. Bulgarelli, D., Rott, M., Schlaeppi, K., Ver Loren van Themaat, E., Ahmadinejad, N., Assenza, F., Rauf, P., Huettel, B., Reinhardt, R., Schmelzer, E., Peplies, J., Gloeckner, F. O., Amann, R., Eickhorst, T. and Schulze-Lefert, P. (2012) ‘Revealing structure and assembly cues for *Arabidopsis* root-inhabiting bacterial microbiota’, Nature. Nature Publishing Group, 488(7409), pp. 91–95. doi: 10.1038/nature11336.

15. Bulgarelli, D., Schlaeppi, K., Spaepen, S., van Themaat, E. V. L. and Schulze-Lefert, P. (2013) ‘Structure and functions of the bacterial microbiota of plants’, Annual Review of Plant Biology, 64(1), pp. 807–838. doi: 10.1146/annurev-arplant-050312-120106.

16. Bulgarelli, D., Garrido-Oter, R., Münch, P. C., Weiman, A., Dröge, J., Pan, Y., McHardy, A. C. and Schulze-Lefert, P. (2015) ‘Structure and function of the bacterial root microbiota in wild and domesticated barley’, Cell Host and Microbe, 17(3), pp. 392–403. doi: 10.1016/j.chom.2015.01.011.

17. Canarini, A., Kaiser, C., Merchant, A., Richter, A. and Wanek, W. (2019) ‘Root exudation of primary metabolites: Mechanisms and their roles in plant responses to environmental stimuli’, Frontiers in Plant Science, 10. doi: 10.3389/fpls.2019.00157.

18. Caporaso, J. G., Kuczynski, J., Stombaugh, J., Bittinger, K., Bushman, F. D., Costello, E. K., Fierer, N., Pẽa, A. G., Goodrich, J. K., Gordon, J. I., Huttley, G. A., Kelley, S. T., Knights, D., Koenig, J. E., Ley, R. E., Lozupone, C. A., McDonald, D., Muegge, B. D., Pirrung, M., Reeder, J., Sevinsky, J. R., Turnbaugh, P. J., Walters, W. A., Widmann, J., Yatsunenko, T., Zaneveld, J. and Knight, R. (2010) ‘QIIME allows analysis of high-throughput community sequencing data’, Nature Methods. Nature Publishing Group, 7(5), pp. 335–336. doi: 10.1038/nmeth.f.303.

19. Chen, H., Jogler, M., Rohde, M., Klenk, H. P., Busse, H. J., Tindall, B. J., Spröer, C. and Overmann, J. (2013) ‘*Sphingobium limneticum* sp. nov. and *Sphingobium boeckii* sp. nov., two freshwater planktonic members of the family *Sphingomonadaceae*, and reclassification of *Sphingomonas suberifaciens* as *Sphingobium suberifaciens* comb. nov’, International Journal of Systematic and Evolutionary Microbiology, 63(2), pp. 735–743. doi: 10.1099/ijs.0.040105-0.

20. Compant, S., Clément, C. and Sessitsch, A. (2010) ‘Plant growth-promoting bacteria in the rhizo- and endosphere of plants: Their role, colonization, mechanisms involved and prospects for utilization’, Soil Biology and Biochemistry, 42(5), pp. 669–678. doi: 10.1016/j.soilbio.2009.11.024.

21. Conway, J. R., Lex, A. and Gehlenborg, N. (2017) ‘UpSetR: an R package for the visualization of intersecting sets and their properties’, Bioinformatics. Edited by J. Hancock, 33(18), pp. 2938–2940. doi: 10.1093/bioinformatics/btx364.

22. Cruz-Ramos, H., Glaser, P., Wray, L. V. and Fisher, S. H. (1997) ‘The *Bacillus subtilis ure*ABC operon.’, Journal of bacteriology, 179(10), pp. 3371–3373. doi: 10.1128/JB.179.10.3371-3373.1997.

23. Delmont, T. O. and Eren, E. M. (2018) ‘Linking pangenomes and metagenomes: The *Prochlorococcus* metapangenome’, PeerJ, 2018(1), p. e4320. doi: 10.7717/peerj.4320.

24. DeSantis, T. Z., Hugenholtz, P., Larsen, N., Rojas, M., Brodie, E. L., Keller, K., Huber, T., Dalevi, D., Hu, P. and Andersen, G. L. (2006) ‘Greengenes, a chimera-checked 16S rRNA gene database and workbench compatible with ARB’, Applied and Environmental Microbiology, 72(7), pp. 5069–5072. doi: 10.1128/AEM.03006-05.

25. de Weert, S., Vermeiren, H., Mulders, I. H. M., Kuiper, I., Hendrickx, N., Bloemberg, G. V., Vanderleyden, J., De Mot, R. and Lugtenberg, B. J. J. (2002) ‘Flagella-driven chemotaxis towards exudate components is an important trait for tomato root colonization by *Pseudomonas fluorescens*’, Molecular Plant-Microbe Interactions, 15(11), pp. 1173–1180. doi: 10.1094/MPMI.2002.15.11.1173.

26. Doebley, J. F., Gaut, B. S. and Smith, B. D. (2006) ‘The molecular genetics of crop domestication’, Cell, pp. 157. doi: 10.1016/j.cell.2006.12.006.

27. Edwards, J., Johnson, C., Santos-Medellín, C., Lurie, E., Podishetty, N. K., Bhatnagar, S., Eisen, J. A., Sundaresan, V. and Jeffery, L. D. (2015) ‘Structure, variation, and assembly of the root-associated microbiomes of rice’, Proceedings of the National Academy of Sciences of the United States of America, 112(8), pp. E911–E920. doi: 10.1073/pnas.1414592112.

28. El-Tarabily, K. A., Nassar, A. H., Hardy, G. E. S. J. and Sivasithamparam, K. (2009) ‘Plant growth promotion and biological control of *Pythium aphanidermatum*, a pathogen of cucumber, by endophytic actinomycetes’, Journal of Applied Microbiology, 106(1), pp. 13–26. doi: 10.1111/j.1365-2672.2008.03926.x.

29. Fierer, N. and Jackson, R. B. (2006) ‘The diversity and biogeography of soil bacterial communities’, Proceedings of the National Academy of Sciences of the United States of America, 103(3), pp. 626–631. doi: 10.1073/pnas.0507535103.

30. Finck-Barbançon, V., Goranson, J., Zhu, L., Sawa, T., Wiener-Kronish, J. P., Fleiszig, S. M. J., Wu, C., Mende-Mueller, L. and Frank, D. W. (1997) ‘ExoU expression by *Pseudomonas aeruginosa* correlates with acute cytotoxicity and epithelial injury’, Molecular Microbiology, 25(3), pp. 547–557. doi: 10.1046/j.1365-2958.1997.4891851.x.

31. Fitzpatrick, C. R., Copeland, J., Wang, P. W., Guttman, D. S., Kotanen, P. M. and Johnson, M. T. J. (2018) ‘Assembly and ecological function of the root microbiome across angiosperm plant species’, Proceedings of the National Academy of Sciences of the United States of America, 115(6), pp. E1157–E1165. doi: 10.1073/pnas.1717617115.

32. Fowler, J. H., Narváez-Vásquez, J., Aromdee, D. N., Pautot, V., Holzer, F. M. and Walling, L. L. (2009) ‘Leucine aminopeptidase regulates defense and wound signaling in tomato downstream of jasmonic acid’, The Plant Cell, 21(4), pp. 1239–1251. doi: 10.1105/tpc.108.065029.

33. Hartmann, A., Rothballer, M. and Schmid, M. (2008) ‘Lorenz Hiltner, a pioneer in rhizosphere microbial ecology and soil bacteriology research’, Plant and Soil, 312(1–2), pp. 7–14. doi: 10.1007/s11104-007-9514-z.

34. Huffman, E. W. D. (1977) ‘Performance of a new automatic carbon dioxide coulometer’, Microchemical Journal, 22(4), pp. 567–573. doi: 10.1016/0026-265X(77)90128-X.

35. Hyatt, D., Chen, G. L., LoCascio, P. F., Land, M. L., Larimer, F. W. and Hauser, L. J. (2010) ‘Prodigal: Prokaryotic gene recognition and translation initiation site identification’, BMC Bioinformatics, 11(1), p. 119. doi: 10.1186/1471-2105-11-119.

36. IUSS Working Group WRB. 2015. World Reference Base for Soil Resources 2014, update 2015. ‘International soil classification system for naming soils and creating legends for soil maps’. World Soil Resources Reports No. 106. FAO, Rome.

37. Inceoǧlu, Ö., Al-Soud, W. A., Salles, J. F., Semenov, A. V. and van Elsas, J. D. (2011) ‘Comparative analysis of bacterial communities in a potato field as determined by pyrosequencing’, PLoS ONE. 6(8), p. e23321. doi: 10.1371/journal.pone.0023321.

38. INEGI, 2014. Conjunto de datos vectoriales edafológico, escala 1:250000 Serie II. (Continuo Nacional), escala: 1:250000. edición: 2. Instituto Nacional de Estadística y Geografía. Aguascalientes, Ags., México.

39. Lang, R., 1920. Verwitterung und bodenbildung als einfuehrung in die bodenkunde. Schweizerbart Science Publishers, Stuttgart.

40. Langmead, B. and Salzberg, S. L. (2012) ‘Fast gapped-read alignment with Bowtie 2’, Nature Methods, 9(4), pp. 357–359. doi: 10.1038/nmeth.1923.

41. Larousse, M., Rancurel, C., Syska, C., Palero, F., Etienne, C., Industri, B., Nesme, X., Bardin, M. and Galiana, E. (2017) ‘Tomato root microbiota and *Phytophthora parasitica*-associated disease’, *Microbiome*. Microbiome, 5(1), p. 56. doi: 10.1186/S40168-017-0273-7.

42. Lauber, C. L., Hamady, M., Knight, R. and Fierer, N. (2009) ‘Pyrosequencing-based assessment of soil pH as a predictor of soil bacterial community structure at the continental scale’, Applied and Environmental Microbiology, 75(15), pp. 5111–5120. doi: 10.1128/AEM.00335-09.

43. Lee, S. A., Park, J., Chu, B., Kim, J. M., Joa, J. H., Sang, M. K., Song, J. and Weon, H. Y. (2016) ‘Comparative analysis of bacterial diversity in the rhizosphere of tomato by culture-dependent and -independent approaches’, Journal of Microbiology, 54(12), pp. 823–831. doi: 10.1007/s12275-016-6410-3.

44. Li, W. and Godzik, A. (2006) ‘Cd-hit: A fast program for clustering and comparing large sets of protein or nucleotide sequences’, Bioinformatics, 22(13), pp. 1658–1659. doi: 10.1093/bioinformatics/btl158.

45. Liu, R. and Ochman, H. (2007) ‘Origins of flagellar gene: Operons and secondary flagellar systems’, Journal of Bacteriology, 189(19), pp. 7098–7104. doi: 10.1128/JB.00643-07.

46. Love, M. I., Huber, W. and Anders, S. (2014) ‘Moderated estimation of fold change and dispersion for RNA-seq data with DESeq2’, Genome Biology, 15(12), p. 550. doi: 10.1186/s13059-014-0550-8.

47. Lozupone, C., Hamady, M. and Knight, R. (2006) ‘UniFrac - An online tool for comparing microbial community diversity in a phylogenetic context’, BMC Bioinformatics, 7(1), p. 371. doi: 10.1186/1471-2105-7-371.

48. Lundberg, D. S., Lebeis, S. L., Paredes, S. H., Yourstone, S., Gehring, J., Malfatti, S., Tremblay, J., Engelbrektson, A., Kunin, V., Rio, T. G. del Edgar, R. C., Eickhorst, T., Ley, R. E., Hugenholtz, P., Tringe, S. G. and Dangl, J. L. (2012) ‘Defining the core *Arabidopsis thaliana* root microbiome’, Nature. Nature Publishing Group, 488(7409), pp. 86–90. doi: 10.1038/nature11237.

49. Ma, P., Patching, S. G., Ivanova, E., Baldwin, J. M., Sharples, D., Baldwin, S. A. and Henderson, P. J. F. (2016) ‘Allantoin transport protein, PucI, from *Bacillus subtilis*: evolutionary relationships, amplified expression, activity and specificity’, Microbiology, 162(5), pp. 823–836. doi: 10.1099/mic.0.000266.

50. Männistö, M. K., Tiirola, M. and Häggblom, M. M. (2007) ‘Bacterial communities in arctic fields of Finnish lapland are stable but highly pH-dependent’, FEMS Microbiology Ecology, 59(2), pp. 452–465. doi: 10.1111/j.1574-6941.2006.00232.x.

51. Marschner, P., Crowley, D. and Yang, C. H. (2004) ‘Development of specific rhizosphere bacterial communities in relation to plant species, nutrition and soil type’, Plant and Soil, 261(1–2), pp. 199–208. doi: 10.1023/B:PLSO.0000035569.80747.c5.

52. Masella, A. P., Bartram, A. K., Truszkowski, J. M., Brown, D. G. and Neufeld, J. D. (2012) ‘PANDAseq: Paired-end assembler for illumina sequences’, BMC Bioinformatics. BioMed Central Ltd, 13(1), p. 31. doi: 10.1186/1471-2105-13-31.

53. McMurdie, P. J. and Holmes, S. (2013) ‘Phyloseq: An R Package for reproducible interactive analysis and graphics of microbiome census data’, PLoS ONE. Edited by M. Watson, 8(4), p. e61217. doi: 10.1371/journal.pone.0061217.

54. Menzel, P., Ng, K. L. and Krogh, A. (2016) ‘Fast and sensitive taxonomic classification for metagenomics with Kaiju’, Nature Communications. Nature Publishing Group, 7(1), p. 11257. doi: 10.1038/ncomms11257.

55. Micallef, S. A., Shiaris, M. P. and Colón-Carmona, A. (2009) ‘Influence of *Arabidopsis thaliana* accessions on rhizobacterial communities and natural variation in root exudates’, Journal of Experimental Botany, 60(6), pp. 1729–1742. doi: 10.1093/jxb/erp053.

56. Mira, A., Pushker, R. and Rodríguez-Valera, F. (2006) ‘The Neolithic revolution of bacterial genomes’, Trends in Microbiology, 14(5), pp. 200–206. doi: 10.1016/j.tim.2006.03.001.

57. Murphy, J. and Riley, J. P. (1962) ‘A modified single solution method for the determination of phosphate in natural waters’, Analytica Chimica Acta, 27(C), pp. 31–36. doi: 10.1016/S0003-2670(00)88444-5.

58. Nawrocki, E. P., Kolbe, D. L. and Eddy, S. R. (2009) ‘Infernal 1.0: Inference of RNA alignments’, Bioinformatics, 25(10), pp. 1335–1337. doi: 10.1093/bioinformatics/btp157.

59. Naylor, D., Degraaf, S., Purdom, E. and Coleman-Derr, D. (2017) ‘Drought and host selection influence bacterial community dynamics in the grass root microbiome’, ISME Journal. Nature Publishing Group, 11(12), pp. 2691–2704. doi: 10.1038/ismej.2017.118.

60. Neilson, J. W., Califf, K., Cardona, C., Copeland, A., van Treuren, W., Josephson, K. L., Knight, R., Gilbert, J. A., Quade, J., Caporaso, J. G. and Maier, R. M. (2017) ‘Significant impacts of increasing aridity on the arid soil microbiome’, mSystems. Edited by H. Chu, 2(3), pp. 1–15. doi: 10.1128/msystems.00195-16.

61. Nurk, S., Meleshko, D., Korobeynikov, A. and Pevzner, P. A. (2017) ‘MetaSPAdes: A new versatile metagenomic assembler’, Genome Research, 27(5), pp. 824–834. doi: 10.1101/gr.213959.116.

62. Oksanen J, Blanchet FG, Kindt R et al. Vegan: Community ecology package. R package version 2.3–0, 2015, https://cran.r-project.org/web/packages/vegan/index.html.

63. Overbeek, R., Olson, R., Pusch, G. D., Olsen, G. J., Davis, J. J., Disz, T., Edwards, R. A., Gerdes, S., Parrello, B., Shukla, M., Vonstein, V., Wattam, A. R., Xia, F. and Stevens, R. (2014) ‘The SEED and the Rapid Annotation of microbial genomes using Subsystems Technology (RAST)’, Nucleic Acids Research, 42(D1), pp. 206–214. doi: 10.1093/nar/gkt1226.

64. Panke-Buisse, K., Poole, A. C., Goodrich, J. K., Ley, R. E. and Kao-Kniffin, J. (2015) ‘Selection on soil microbiomes reveals reproducible impacts on plant function’, ISME Journal. Nature Publishing Group, 9(4), pp. 980–989. doi: 10.1038/ismej.2014.196.

65. Pautot, V., Holzer, F. M., Chaufaux, J. and Walling, L. L. (2001) ‘The induction of tomato leucine aminopeptidase genes (LapA) after *Pseudomonas syringae* pv. tomato infection is primarily a wound response triggered by coronatine’, Molecular Plant-Microbe Interactions, 14(2), pp. 214–224. doi: 10.1094/MPMI.2001.14.2.214.

66. Pautot, V., Holzer, F. M., Reisch, B. and Walling, L. L. (1993) ‘Leucine aminopeptidase: an inducible component of the defense response in *Lycopersicon esculentum* (tomato).’, Proceedings of the National Academy of Sciences, 90(21), pp. 9906–9910. doi: 10.1073/pnas.90.21.9906.

67. Peiffer, J. A., Spor, A., Koren, O., Jin, Z., Tringe, S. G., Dangl, J. L., Buckler, E. S. and Ley, R. E. (2013) ‘Diversity and heritability of the maize rhizosphere microbiome under field conditions’, Proceedings of the National Academy of Sciences of the United States of America, 110(16), pp. 6548–6553. doi: 10.1073/pnas.1302837110.

68. Pérez-Jaramillo, J. E., Carrión, V. J., de Hollander, M. and Raaijmakers, J. M. (2018) ‘The wild side of plant microbiomes’, *Microbiome*. Microbiome, 6(1), p. 143. doi: 10.1186/s40168-018-0519-z.

69. Phillips, R. M., Six, D. A., Dennis, E. A. and Ghosh, P. (2003) ‘In vivo phospholipase activity of the *Pseudomonas aeruginosa* cytotoxin ExoU and protection of mammalian cells with phospholipase A 2 inhibitors’, Journal of Biological Chemistry, 278(42), pp. 41326–41332. doi: 10.1074/jbc.M302472200.

70. Pii, Y., Borruso, L., Brusetti, L., Crecchio, C., Cesco, S. and Mimmo, T. (2016) ‘The interaction between iron nutrition, plant species and soil type shapes the rhizosphere microbiome’, Plant Physiology and Biochemistry. Elsevier Masson SAS, 99, pp. 39–48. doi: 10.1016/j.plaphy.2015.12.002.

71. Poindexter, J. S. (1981) ‘The Caulobacters: Ubiquitous unusual bacteria’, Microbiological Reviews, pp. 123–179. doi: 10.1128/mmbr.45.1.123-179.1981.

72. Price, M. N., Dehal, P. S. and Arkin, A. P. (2009) ‘Fasttree: Computing large minimum evolution trees with profiles instead of a distance matrix’, Molecular Biology and Evolution, 26(7), pp. 1641–1650. doi: 10.1093/molbev/msp077.

73. Pruitt, K. D., Tatusova, T. and Maglott, D. R. (2007) ‘NCBI reference sequences (RefSeq): A curated non-redundant sequence database of genomes, transcripts and proteins’, Nucleic Acids Research, 35(SUPPL. 1), pp. D61–D65. doi: 10.1093/nar/gkl842.

74. Purugganan, M. D. and Fuller, D. Q. (2009) ‘The nature of selection during plant domestication’, Nature, pp. 843–848. doi: 10.1038/nature07895.

75. Ramin, K. I. and Allison, S. D. (2019) ‘Bacterial tradeoffs in growth rate and extracellular enzymes’, Frontiers in Microbiology, 10, pp. 1–10. doi: 10.3389/fmicb.2019.02956.

76. Redford, A. J., Bowers, R. M., Knight, R., Linhart, Y. and Fierer, N. (2010) ‘The ecology of the phyllosphere: Geographic and phylogenetic variability in the distribution of bacteria on tree leaves’, Environmental Microbiology, 12(11), pp. 2885–2893. doi: 10.1111/j.1462-2920.2010.02258.x.

77. Roesch, L. F. W., Fulthorpe, R. R., Riva, A., Casella, G., Hadwin, A. K. M., Kent, A. D., Daroub, S. H., Camargo, F. A. O., Farmerie, W. G. and Triplett, E. W. (2007) ‘Pyrosequencing enumerates and contrasts soil microbial diversity’, ISME Journal, 1(4), pp. 283–290. doi: 10.1038/ismej.2007.53.

78. Sato, H. (2003) ‘The mechanism of action of the *Pseudomonas aeruginosa*-encoded type III cytotoxin, ExoU’, The EMBO Journal, 22(12), pp. 2959–2969. doi: 10.1093/emboj/cdg290.

79. Satour, M. M., and E. E. Butler. “A root and crown rot of tomato caused by *Phytophthora capsici* and *P. parasitica*.” Phytopathology 57, no. 5 (1967): 510.

80. Scheller, H. V. and Ulvskov, P. (2010) ‘Hemicelluloses’, Annual Review of Plant Biology, 61(1), pp. 263–289. doi: 10.1146/annurev-arplant-042809-112315.

81. Schlaeppi, K., Dombrowski, N., Oter, R. G., Ver Loren Van Themaat, E. and Schulze-Lefert, P. (2014) ‘Quantitative divergence of the bacterial root microbiota in *Arabidopsis thaliana* relatives’, Proceedings of the National Academy of Sciences of the United States of America, 111(2), pp. 585–592. doi: 10.1073/pnas.1321597111.

82. Schreiter, S., Ding, G. C., Heuer, H., Neumann, G., Sandmann, M., Grosch, R., Kropf, S. and Smalla, K. (2014) ‘Effect of the soil type on the microbiome in the rhizosphere of field-grown lettuce’, Frontiers in Microbiology, 5, p. 144. doi: 10.3389/fmicb.2014.00144.

83. Skerker, J. M. and Shapiro, L. (2000) ‘Identification and cell cycle control of a novel pilus system in *Caulobacter crescentus*’, The EMBO Journal, 19(13), pp. 3223–3234. doi: 10.1093/emboj/19.13.3223.

84. Sreevidya, M., Gopalakrishnan, S., Kudapa, H. and Varshney, R. K. (2016) ‘Exploring plant growth-promotion actinomycetes from vermicompost and rhizosphere soil for yield enhancement in chickpea’, Brazilian Journal of Microbiology. Sociedade Brasileira de Microbiologia, 47(1), pp. 85–95. doi: 10.1016/j.bjm.2015.11.030.

85. Swenson, W., Wilson, D. S. and Elias, R. (2000) ‘Artificial ecosystem selection’, Proceedings of the National Academy of Sciences of the United States of America, 97(16), pp. 9110–9114. doi: 10.1073/pnas.150237597.

86. van Der Heijden, M. G. A., Bardgett, R. D. and Van Straalen, N. M. (2008) ‘The unseen majority: Soil microbes as drivers of plant diversity and productivity in terrestrial ecosystems’, Ecology Letters, 11(3), pp. 296–310. doi: 10.1111/j.1461-0248.2007.01139.x.

87. Thompson, Luke R., Jon G. Sanders, Daniel McDonald, Amnon Amir, Joshua Ladau, Kenneth J. Locey, Robert J. Prill, et al. 2017. “A communal catalogue reveals Earth’s multiscale microbial diversity.” Nature 551 (7681): 457–63. https://doi.org/10.1038/nature24621.

88. Tian, B. Y., Cao, Y. and Zhang, K. Q. (2015) ‘Metagenomic insights into communities, functions of endophytes, and their associates with infection by root-knot nematode, *Meloidogyne incognita*, in tomato roots’, Scientific Reports. Nature Publishing Group, 5(1), p. 17087. doi: 10.1038/srep17087.

89. Watzer, B. and Forchhammer, K. (2018) ‘Cyanophycin synthesis optimizes nitrogen utilization in the unicellular cyanobacterium *Synechocystis* sp. strain PCC 6803’, Applied and Environmental Microbiology. Edited by C. Vieille, 84(20), pp. 1–18. doi: 10.1128/AEM.01298-18.

90. Wilke, A., Harrison, T., Wilkening, J., Field, D., Glass, E. M., Kyrpides, N., Mavrommatis, K. and Meyer, F. (2012) ‘The M5nr: A novel non-redundant database containing protein sequences and annotations from multiple sources and associated tools’, BMC Bioinformatics, 13(1), p. 141. doi: 10.1186/1471-2105-13-141.

91. Wilkinson, L. (2011) ‘ggplot2: Elegant Graphics for Data Analysis by WICKHAM, H.’, Biometrics, 67(2), pp. 678–679. doi: 10.1111/j.1541-0420.2011.01616.x.

92. Wissuwa, M., Mazzola, M. and Picard, C. (2009) ‘Novel approaches in plant breeding for rhizosphere-related traits’, Plant and Soil, 321(1-2), pp. 409–430. doi: 10.1007/s11104-008-9693-2.

93. Wu, H., Xiang, W., Ouyang, S., Forrester, D. I., Zhou, B., Chen, L., Ge, T., Lei, P., Chen, L., Zeng, Y., Song, X., Peñuelas, J. and Peng, C. (2019) ‘Linkage between tree species richness and soil microbial diversity improves phosphorus bioavailability’, Functional Ecology., 33(8), pp. 1549–1560. doi: 10.1111/1365-2435.13355.

94. Yang, Z., Yang, W., Li, S., Hao, J., Su, Z., Sun, M., Gao, Z. and Zhang, C. (2016) ‘Variation of bacterial community diversity in rhizosphere soil of sole-cropped versus intercropped Wheat field after harvest’, PLoS ONE, 11(3), pp. 1–18. doi: 10.1371/journal.pone.0150618.

95. Young, C. C., Arun, A. B., Kämpfer, P., Busse, H. J., Lai, W. A., Chen, W. M., Shen, F. T. and Rekha, P. D. (2008) ‘*Sphingobium rhizovicinum* sp. nov., isolated from rhizosphere soil of *Fortunella hindsii* (Champ. ex Benth.) Swingle’, International Journal of Systematic and Evolutionary Microbiology, 58(8), pp. 1801–1806. doi: 10.1099/ijs.0.65564-0.

96. Zerbino, D. R. and Birney, E. (2008) ‘Velvet: Algorithms for de novo short read assembly using de Bruijn graphs’, Genome Research, 18(5), pp. 821–829. doi: 10.1101/gr.074492.107.

